# Viscoelastic liquid lassos extend the defensive reach of termite soldiers

**DOI:** 10.64898/2026.10.27.762288

**Authors:** Elio J. Challita, Daniel C. Ames, Jacob S. Harrison, Christina McDonald, Pankaj Rohilla, Prateek Sehgal, David A. Nickle, Jochen Mueller, Saad Bhamla

## Abstract

Small animals face a severe size asymmetry when defending themselves against larger predators. Soldiers of *Nasutitermes* termites overcome this constraint by ejecting tens of nanoliters of secretion through a cephalic nozzle, the nasus. High-speed imaging in the field, rheometry, micro-CT, flow simulation, behavioral assays, and scaled experiments reveal a multistage defense. A near-linear nasus and shear-thinning secretion support rapid ejection. Lateral head oscillation at 5–30 Hz couples with rope-coiling instability to distribute the filament as loops and meanders, a “liquid lasso,” reaching up to three body lengths. After deposition, the apparent extensional viscosity increases 8–80-fold, delaying breakup as the secretion cures into a persistent adhesive. The lasso coats the sensory and grooming structures of predatory ants, and tethers them to the substrate. Synthetic experiments show that nozzle geometry, orientation, height, and oscillatory motion jointly control ejection and coverage, providing design principles for deploying small volumes of time-evolving viscoelastic materials.

## INTRODUCTION

Fluid ejection is widespread in animal defense and predation. Spitting cobras, archerfish, bombardier beetles, velvet worms, spitting spiders (*Scytodes* spp.), and wood ants (*Formica rufa* Linnaeus, 1761) [1] deploy jets, sprays, or filaments to deter or subdue other organisms [2]. Effective deployment, however, imposes competing mechanical demands: the ejected fluid must flow rapidly through a small outlet, remain coherent during flight, and intercept a moving target. These demands are particularly severe for millimeter-scale animals operating through micrometer-scale nozzles with limited fluid reserves.

Soldiers of *Nasutitermes* termites confront these constraints using a resin-rich secretion discharged through a pointed cephalic nozzle, the nasus (Fig. 1a,b; Fig. S15). Although only ∼ 5 mm long, each soldier contains a hypertrophied frontal gland whose secretion can constitute nearly 10% of its body mass and consists primarily of monoterpene solvents, including *α*-pinene and limonene, and diterpenoid resins [3–5]. This defense is particularly important against ants, which are major predators of termites and are frequently larger than the soldiers themselves. The trap-jaw ant in Fig. 1a, for example, is roughly twice the soldier’s length. Ants exploit breaches in nests and shelter structures to attack exposed colony members [6–11]. When the colony is breached (Fig. 2a,b), the largely blind soldiers detect disturbances with their antennae, orient the nasus toward the threat, and eject the secretion (Fig. 1c), leading Eisner to describe them as “ambulatory spray guns” [12]. The secretion acts as an alarm signal, behavioral deterrent, and persistent adhesive [5, 13, 14]. Contaminated ants slow down to termite walking speeds [12] and groom up to 50-fold more after contact than before (Fig. S2; SI Section II), disrupting both movement and continued attack.

**FIG. 1.**
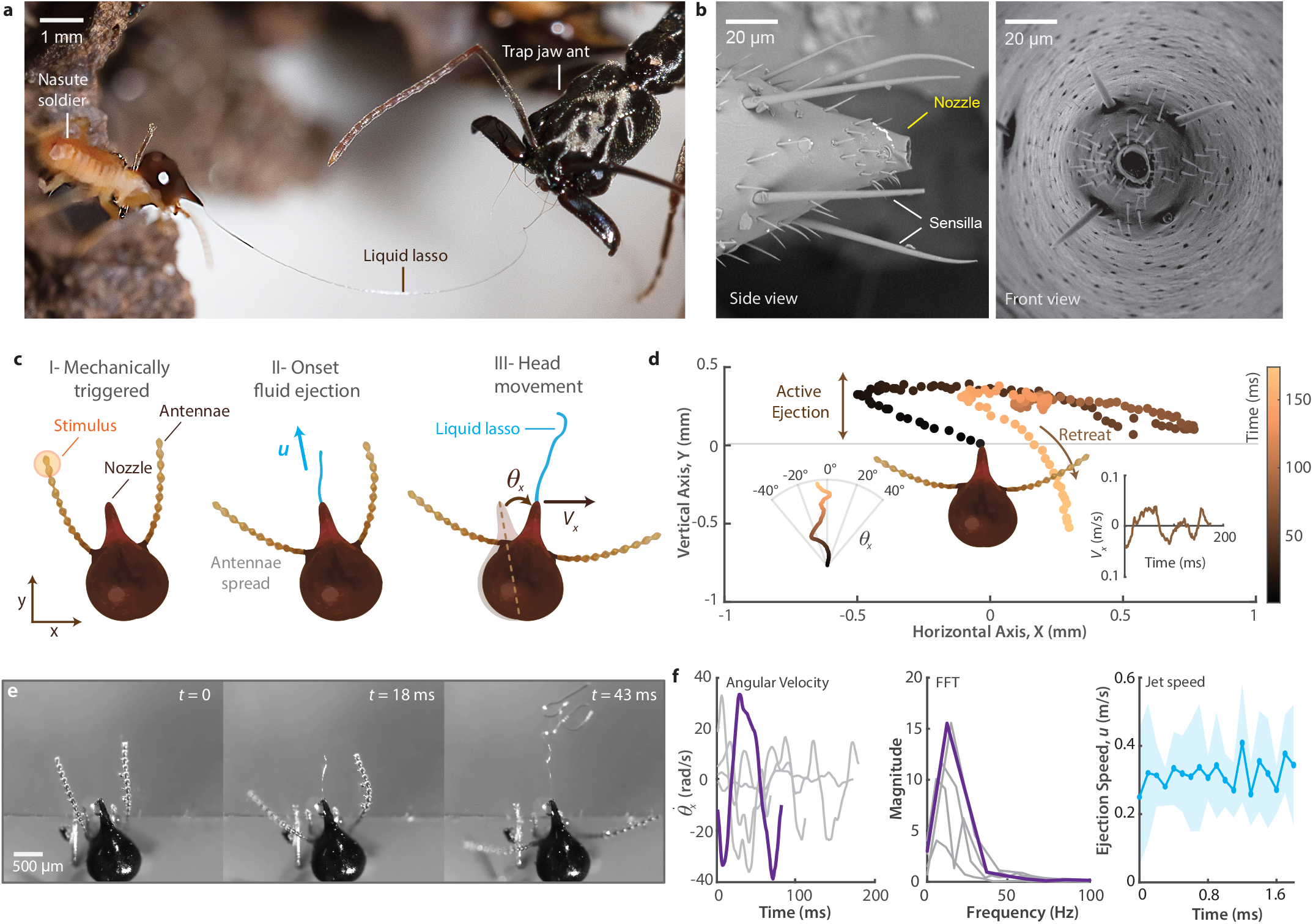
*Nasutitermes* soldiers produce liquid lassos by coupling rapid ejection with lateral head oscillation. **a**. A *Nasutitermes corniger* soldier ejecting viscoelastic secretion toward a trap-jaw ant in the field at Fond Doux, Saint Lucia. **b**. Scanning electron micrographs of the nasus, the cephalic ejection nozzle. Side and front views (*Nasutitermes* sp. 1). The nozzle opening is located at the tip of the head and is surrounded by sensilla. **c**. Schematic of the kinematic measurements following mechanical stimulation of the antennae. We track the jet speed *u* near the nozzle exit, the angular velocity 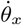 of the head, and the lateral velocity *V*_*x*_ of the head. **d**. Top view of a representative head trajectory during defensive ejection, showing lateral oscillation over an angular excursion of approximately 30^*°*^before the soldier retreats. Peak-to-peak angular excursions ranged from 10^*°*^to 40^*°*^across individuals (*n* = 5). **e**. High-speed image sequence showing the onset of secretion ejection through the nasus onto a glass slide. **f**. Head angular velocity, corresponding frequency spectrum, and jet speed during a representative ejection. Across individuals, head oscillation frequencies ranged from 5 to 30 Hz.

**FIG. 2.**
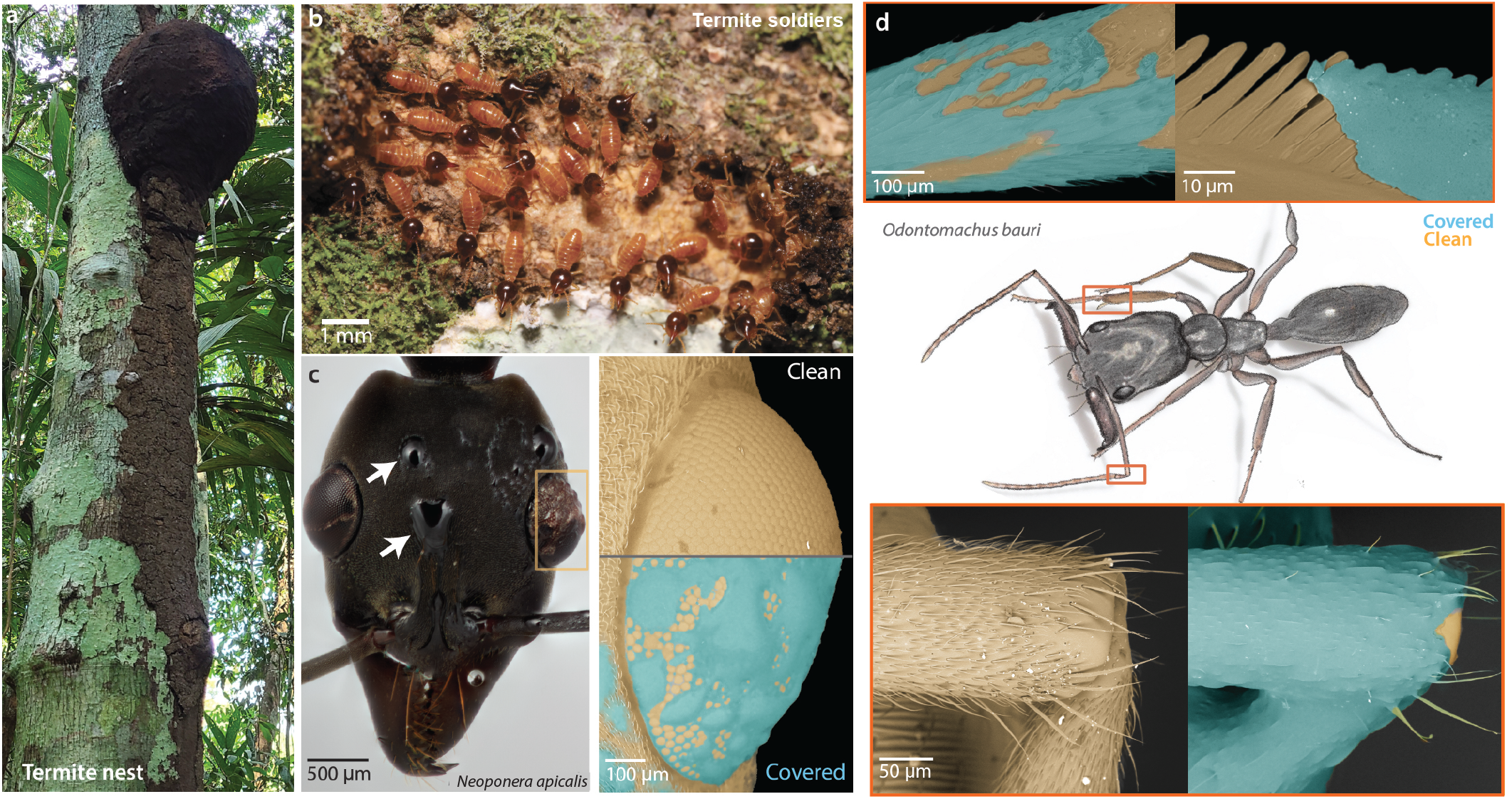
Termite secretion coats the sensory and grooming structures of ant predators. **a**. Arboreal *Nasutitermes* nest on a tree trunk. **b**. Soldiers emerging at a disturbed region of the nest surface. **c**. Head of the termite-hunting ant *Neoponera apicalis* (Latreille, 1802) [18] after encounter with termites, with secretion near the eyes and antennae (arrows). Right: scanning electron micrographs of the compound eye, clean and secretion-covered. **d**. Trap-jaw ant, *Odontomachus bauri* Emery, 1892 [19], after an encounter, with scanning electron micrographs of the foreleg antenna cleaner (strigil) and of the leg surface, each shown covered and clean. In c and d, false color marks secretion-covered (cyan) and clean (orange) cuticle. Scale bars as indicated.

Previous studies have characterized the chemical composition of nasute secretion [4, 5] and qualitatively described its adhesive and immobilizing effects [12]. However, the physical mechanisms that transform a discharge of only tens of nanoliters into an effective defense remain unresolved. How fast and how far can a soldier project such a small volume through an aperture only tens of micrometers wide? How does a narrow, fixed nasus distribute this limited volume across a moving predator larger than the soldier itself without producing a wide spray? How can the same secretion flow readily during ejection yet resist removal once deposited? And are the geometry of the nasus and the motion of the head matched to the rheology of the fluid they deploy? Answering these questions requires linking secretion rheology, nasus morphology, and soldier kinematics across the successive stages of deployment.

Here, we combine high-speed imaging in the field, steady-shear and extensional rheometry, micro-computed tomography, computational fluid dynamics, behavioral assays, and scaled physical experiments to resolve this sequence. We show that nasute termite defense operates as a multistage fluid-mechanical system. A near-linearly tapered nasus and shear-thinning secretion reduce resistance during rapid extrusion, while the ejected filament remains coherent over body-scale distances. Lateral head oscillation sweeps the narrow jet across the target, and substrate-induced rope coiling deposits the filament as overlapping loops, meanders, and bridges, forming a network that we term a “liquid lasso.” Following contact, extensional strain hardening delays filament breakup, while aging progressively converts the secretion into a persistent, solid-like adhesive. During ant encounters, the secretion adheres to sensory and grooming structures, elicits increased grooming, and forms mechanical tethers between the predator and substrate. Finally, experiments with a rheologically informed synthetic analog reveal how nozzle geometry, imposed motion, and evolving material properties jointly govern the ejection and deposition of viscoelastic filaments.

## RESULTS

### Soldiers eject nanoliter volumes as coherent, high-speed threads

Using high-speed imaging, we characterized defensive ejection by arboreal *Nasutitermes* soldiers, including *Nasutitermes corniger* (Motschulsky, 1855) [15] and unidentified *Nasutitermes* sp., collected from the Peruvian Amazon, Barro Colorado Island, Panama, and Costa Rica (SI Section I). Mechanical stimulation of the antennae reliably triggered ejection. Before discharging the secretion, soldiers spread their antennae laterally and oriented the nasus toward the stimulus (Fig. 1c).

We measured the fluid speed *u* near the nozzle exit for *Nasutitermes* sp. before the jet contacted the substrate. The jet diameter measured from high-speed images was *d*_*j*_ = 35 ± 10 *µ*m. Across *n* = 11 individuals, the secretion was expelled at a mean exit speed of *u*_*o*_ = 0.32 ± 0.15 m/s and a mean per-individual peak speed of *u*_max_ = 0.65 ± 0.23 m/s (mean ± SD), with an ejection duration of *τ* = 125 ± 52 ms (Fig. S1). Approximating the ejected filament as a cylindrical jet gives an estimated volume per event of 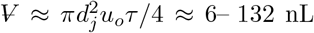. This range is consistent with classical squeeze-extraction measurements from *Nasutitermes exitiosus* (Hill, 1925) [16] soldiers, which yielded a mean secretion volume of 67 ± 9 nL, a maximum of 92 nL, and ejection durations no longer than 105 ms [12].

Soldiers frequently discharged multiple times, indicating that a single ejection does not necessarily exhaust the frontal gland. The mean jet diameter also exceeds that of the aperture, which measures ∼14–32 *µ*m by SEM and micro-CT (Fig. 1b; SI Section IV), consistent with die swell of a viscoelastic extrudate whose stored elastic stress relaxes on exit [17]. Once ejected, the filament travels as a single coherent thread rather than a spray, as expected when viscous and elastic stresses dominate surface tension at this scale (SI Section III). Despite emerging as a single thread only tens of micrometers wide, the secretion reached targets up to three soldier body lengths from the nasus and was deposited over a region far broader than the thread itself (Supplementary video S1). We next examine the two mechanisms that produce this coverage.

### Head oscillation and rope coiling amplify spatial coverage

As the termite secretion reaches a surface, it forms loops and coils characteristic of liquid rope coiling, a classical instability observed when a viscous filament falls onto a substrate [20–22] (Supplementary video S2). In this phenomenon, the filament buckles under axial compression near the point of contact, generating loops whose frequency and size depend on the nozzle–substrate distance, the ejection speed, and the relative importance of viscous, inertial, and gravitational forces.

In the viscous regime of rope coiling, the coiling frequency is expected to scale as Ω ∼ *πu/H*, where *u* is the filament speed and *H* is the nozzle–substrate distance, while the coil diameter is expected to scale approximately with *H* [21, 22]. Across nozzle–substrate distances of 0.1 to 1 mm, we measured coiling angular frequencies ranging from approximately 250 to 3000 Hz (Fig. 3a,b). Fitting the measured frequency–height relation yielded characteristic filament speeds of *u* ∼ 0.03 to 0.24 m*/*s across individuals (*n* = 8, *Nasutitermes* sp.), with a representative fitted value of *u* ∼ 0.26 m*/*s. An independent estimate based on deposited coil kinematics, *u* ∼ *d*Ω, gave a comparable value of 0.12 ± 0.07 m*/*s. Coil diameter also increased approximately linearly with nozzle–substrate distance, *d* = 0.26*H* (*R*^2^ = 0.8, intercept of order 10^*−*5^ m). These measurements support the interpretation that the deposited filament coils in the viscous regime.

**FIG. 3.**
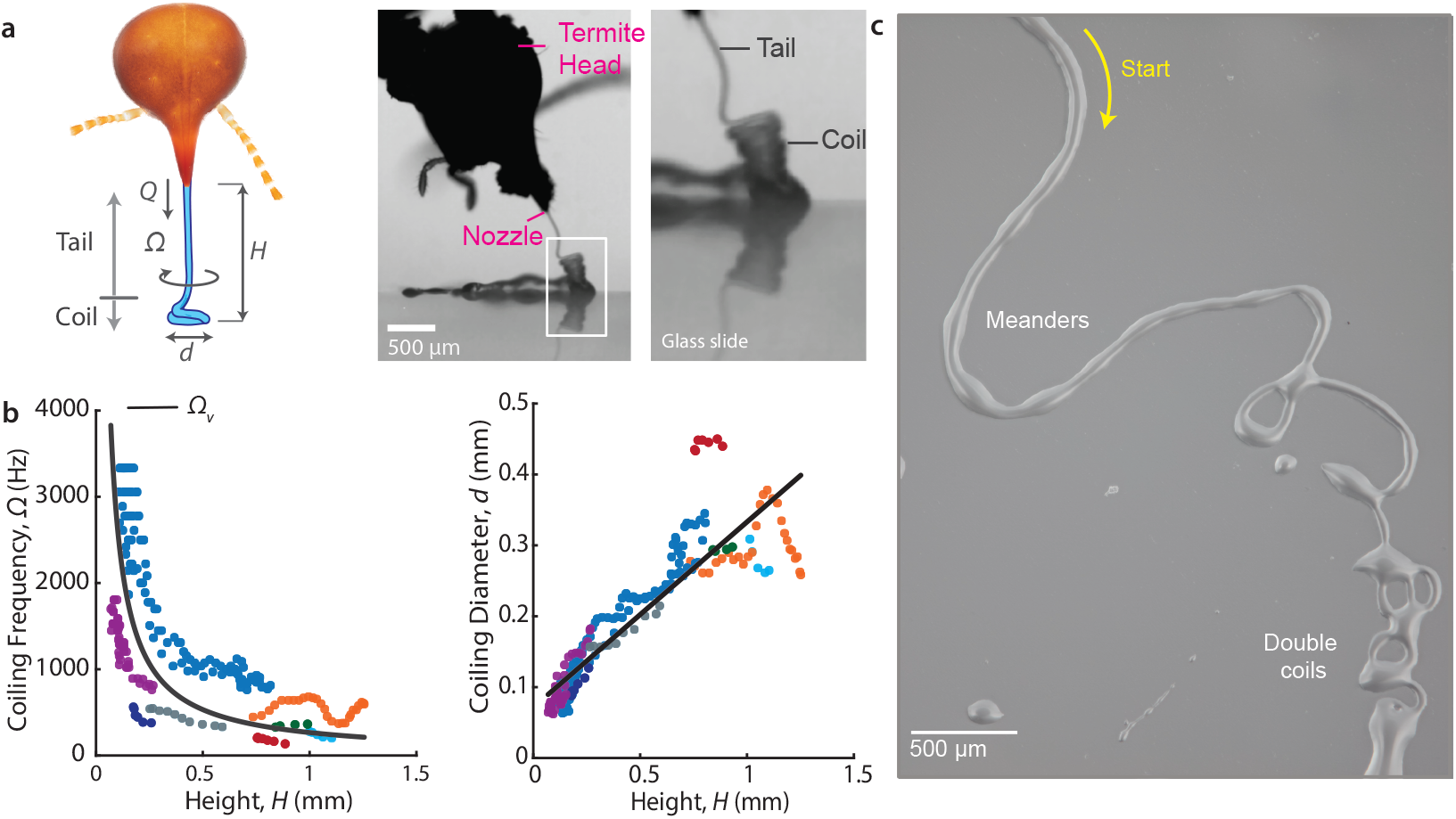
Head scanning and rope coiling spread a single thread into loops, meanders, and double coils. **a**. High-speed imaging of a viscoelastic filament ejected from a termite nasus and coiling upon contact with a glass slide. The schematic defines the nozzle–substrate distance *H*, coiling frequency Ω, coil diameter *d*, and flow rate *Q*. **b**. Coiling frequency decreases with increasing nozzle–substrate distance, while coil diameter increases approximately linearly with distance, consistent with coiling in the viscous regime. Different colors correspond to different individuals of *N*. sp. The black curve is the viscous-regime prediction Ω = *πu/H*. **c**. Lateral head motion during ejection causes the deposited filament to transition from simple coils to more complex patterns, including meanders and double coils.

In the natural defensive behavior, however, the nozzle is not stationary. During ejection, the soldier swept its head laterally in approximately sinusoidal arcs, scanning the filament across the target (Fig. 1d–f). The mean lateral head speed was *V*_*x*_ = 2*V*_*x*,max_*/π* = 0.05 ± 0.01 m*/*s, with a peak speed of *V*_*x*,max_ = 0.077 ± 0.015 m/s across *n* = 5 individuals. The head oscillated at 5–30 Hz with peak-to-peak angular excursions of approximately 10^*°*^– 40^*°*^, producing a lateral sweep of approximately 1.25 mm perpendicular to the initial ejection direction. Eisner et al. previously noted that the head of *N. exitiosus* “vibrates from side to side” during ejection [12]; here we resolve the speed, frequency, and angular excursion of this motion. Following each ejection pulse, the soldier typically retreated.

The oscillation frequency lies within the range of synchronous, muscle-driven insect motion [23]. This active scanning differs from the ∼ 30–60 Hz oscillation of velvet-worm oral papillae, which arises passively through an elastohydrodynamic instability of the slime jet [24]; comparable scanning also occurs in spitting spiders [25].

With the nozzle sweeping and the target itself in motion, the secretion deposited as a dynamically patterned filament rather than as a stationary coil. The ratio of filament speed to lateral head speed, *u/V*_*x*_, spanned approximately 0.6 to 4.8, giving rise to a range of deposited motifs, from simple coils to double coils and meanders (Fig. 3c; Fig. S3) [26, 27]. These patterns closely resemble those produced by a viscous filament falling onto a translating surface, often referred to as a “liquid sewing machine” [28]. Unlike the classical liquid sewing machine, however, the termite nozzle does not translate steadily in one direction. Instead, the head undergoes reciprocal scanning: it sweeps laterally, decelerates, reverses direction, and then retreats. This reciprocal motion causes the deposited filament to pass through multiple patterning regimes during a single ejection event [29].

The lateral sweep of the head acts in flight, carrying the filament sideways and whipping the airborne thread into curls before it lands (Fig. 1e). Axial compression against the substrate acts on contact, driving the rope-coiling instability described above. The sweep thus sets where the filament arrives, coiling how it folds once there, and together they spread the secretion over a footprint that neither alone would produce. A straight, non-coiling jet would remain confined to a narrow line of impact. By contrast, the coiling filament deposits overlapping loops and strands over a much broader region, increasing the probability of contacting multiple body regions of an attacking predator. At the scale of ant–termite interactions, the resulting loops are comparable in size to appendages and cuticular microstructures (Fig. 2d), allowing the secretion to form bridges both across the target and between the target and the substrate. We use the term “liquid lasso” to describe this deposited network of loops, meanders, and bridges produced by the coupling of active head scanning and passive filament instability.

### Strain hardening and aging stabilize the deposited network

The scanning and coiling motions described above distribute the secretion into loops and bridges, but these structures can function as restraints only if they resist breakup when stretched by a moving predator. To characterize the post-ejection mechanical response, we conducted extensional rheology experiments in the field using a custom-built rheometer coupled with high-speed imaging (Methods I). This deformation mode is relevant to defense: as an ant moves, grooms, or attempts to detach, deposited filaments are stretched between appendages, sensory structures, grooming structures (Fig. 2c,d), and the substrate.

Secretions from *N. corniger* and *Nasutitermes* sp. were placed between parallel plates, extended to a final separation *h*_*f*_, and allowed to thin under capillary forces (Fig. 4a,b). We tracked the midplane filament diameter *D*(*t*), normalized by its value *D*_1_ immediately after extension, as a function of time (Fig. 4d; Fig. S4 and SI Section III).

**FIG. 4.**
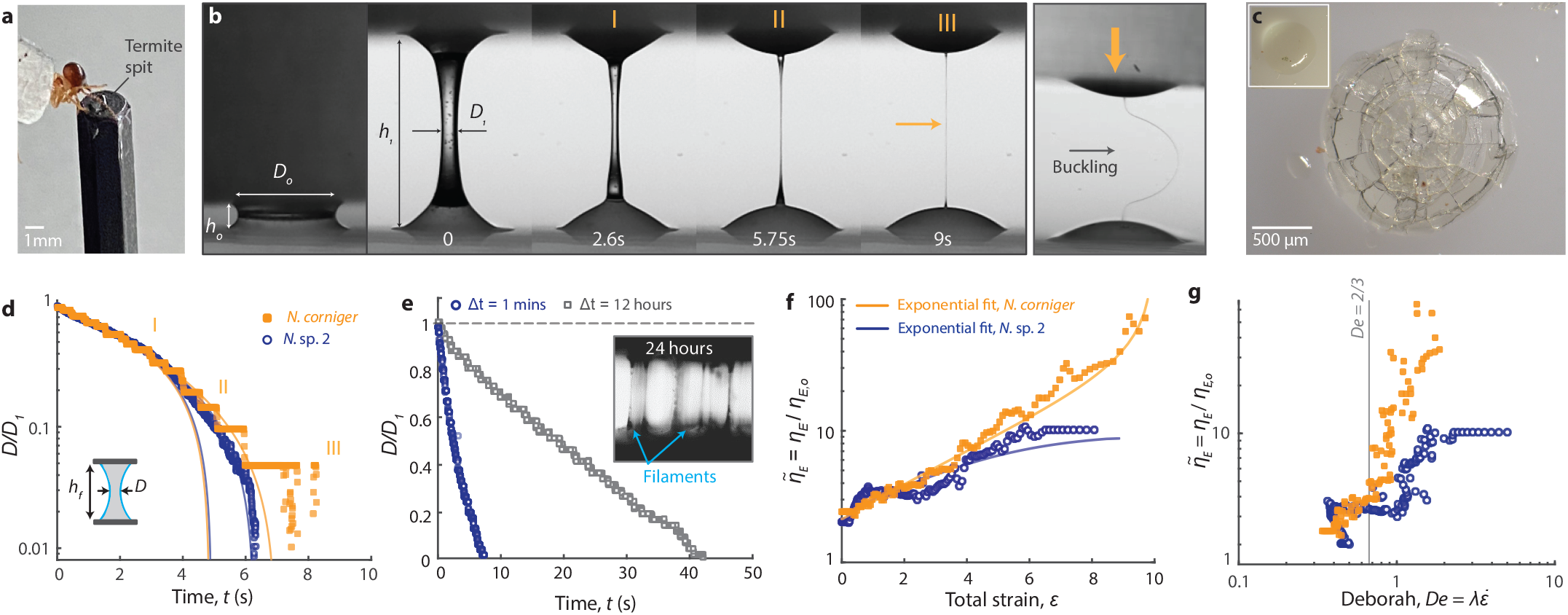
Strain hardening and aging stabilize termite defensive secretion after deposition. **a**. Nasute soldiers were induced to deposit defensive secretion for extensional-rheology measurements. **b**. Capillary thinning of *Nasutitermes corniger* secretion. The initially hourglass-shaped liquid bridge thins into a persistent filament over approximately 9 s. When the upper plate is lowered, the arrested filament buckles as a slender solid-like column. **c**. Secretion collected approximately one year earlier (*Nasutitermes corniger* ), fully solidified and fractured when compressed with a glass slide. Inset: the sample before compression. **d**. Normalized midplane filament diameter *D/D*_1_ as a function of time for *N. corniger* and *Nasutitermes* sp. during capillary thinning. The longer-lived *N. corniger* filaments exhibit a stronger elastic response and delayed breakup. The inset shows the initial bridge geometry. **e**. Filament thinning after different aging intervals. Secretion aged for 12 h exhibits greater filament stability than fresh secretion, while samples aged for 24 h show arrested breakup and a persistent solid-like response (inset). **f**. Normalized apparent extensional viscosity 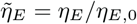 as a function of Hencky strain *ε* [30]. *N. corniger* undergoes pronounced strain hardening, whereas *Nasutitermes* sp. exhibits a weaker increase in extensional viscosity. The early-time viscocapillary baseline, 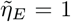, is shown for reference. **g**. Normalized apparent extensional viscosity as a function of Deborah number [31], 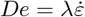. The onset of the elastic-dominated response occurs near *De* = 2*/*3.

The capillary-thinning dynamics exhibited three regimes. During the initial viscocapillary regime (Regime I, *t* ≲ 4 s), viscous stresses balanced surface tension and the filament diameter decreased approximately linearly:

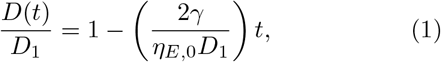

where *γ* is the surface tension and *η*_*E*,0_ is the early-time apparent extensional viscosity. Fits over *t <* 2 s yielded *η*_*E*,0_ ∼ 40 Pa·s for *Nasutitermes* sp. and *η*_*E*,0_ ∼ 200 Pa·s. *corniger*.

During the subsequent elastocapillary regime (Regime II, *t* ≳ 4 s), thinning slowed as elastic stresses developed. The filament diameter followed the expected exponential form [32, 33],

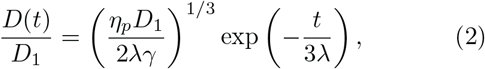

where *η*_*p*_ is the polymeric contribution to the viscosity and *λ* is the extensional relaxation time. Fits to this regime gave *λ* = 1.54 ± 0.88 s (SI Section III).

The two taxa diverged in the terminal regime (Regime III). Filaments from *Nasutitermes* sp. continued thinning until capillary breakup, whereas *N. corniger* filaments approached an arrested, solid-like state and remained intact over the experimental observation period. When the upper plate was lowered, these persistent filaments buckled rather than flowing laterally (Supplementary video S2), demonstrating that the secretion had developed sufficient elasticity or rigidity to behave as a slender column (Fig. 4b).

We quantified strain hardening by plotting the normalized apparent extensional viscosity, 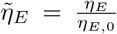 against the accumulated Hencky strain, 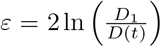. At low strain, 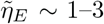, which is consistent with a viscous response [34]. Above a critical strain *ε*_*c*_ ∼ 1–2, 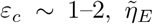 increased, reaching approximately 8–10 for *Nasutitermes* sp. and 70–80 for *N. corniger* (Fig. 4f).

The same transition was evident when 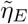 was plotted against the Deborah number, 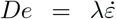, where 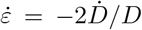 is the instantaneous extension rate. The response transitioned from predominantly viscous to elasticity-dominated near *De* = 2*/*3, above which strain hardening delayed capillary breakup (Fig. 4g) [34, 35]. Extension therefore raises the apparent extensional resistance, allowing deposited bridges to remain intact while a predator moves or grooms. Consistent with this elastic response, filaments deposited on glass (Fig. S3) and on ants (Supplementary video S1) develop a beads-on-a-string morphology, the signature of a viscoelastic thread in which capillary pinching is arrested by extensional stress [36].

The secretion also evolved with time after deposition. After 12 h of aging, the capillary-breakup time increased to approximately 40 s, roughly four times that of fresh secretion, while the apparent extensional viscosity increased to *η*_*E*_ ∼ 3500 Pa·s (Fig. 4e). After 24 h, the aged material no longer underwent capillary breakup under the applied test conditions and instead peeled from the plates like a cured adhesive (Fig. 4e, inset). A separately stored sample collected approximately one year earlier was fully solidified and fractured when compressed with a glass slide Fig. 4c), consistent with the long-term hardening described in early observations of nasute secretion [12].

On an ant predator, the deposited lasso coats sensory and grooming structures. Scanning electron micrographs of ants recovered after termite encounters show the secretion spread across the compound eye of *Neoponera apicalis* (Fig. 2c) and on the foreleg antenna cleaner (strigil) of *Odontomachus bauri* (Fig. 2d). We also conducted field assays in which ants of three taxa (*Eciton harpax, Camponotus* sp., and *Crematogaster* sp.) were filmed for one minute before and one minute after a one-minute encounter with soldier termites. Exposed ants spent up to 50-fold more time grooming after the encounter than before, whereas unexposed controls showed no change. Some ants became tethered to the substrate by filaments of secretion, and material left on the body hardened into a rigid coating overnight (SI Section II, Fig. S2; Supplementary video S1).

### A tapered nasus and shear-thinning rheology enable rapid ejection

So far we have addressed what the secretion does after it leaves the nasus. Ejecting it in the first place imposes a competing requirement: the fluid must pass through an aperture tens of micrometers wide in about a tenth of a second. How readily it does so depends on both its shear rheology and the channel’s shape. To separate these two contributions, we reconstructed the internal profile of the termite nasus from micro-CT data and engineered a scaled extrusion system. Since direct steady-shear characterization of the natural secretion was precluded by the limited available volumes, we modeled its extrusion behavior with Herschel–Bulkley [37] parameters inferred from extensional measurements, ejection kinematics, and a pressure-constrained flow model. We then used a Pluronic-based hydrogel with comparable yield-stress and shear-thinning behavior over the extrusion-relevant shear-rate range as a synthetic analog, which allowed the nozzle geometry and applied pressure to be varied independently without depleting the limited natural secretion.

We reconstructed the internal nasus profile from micro-CT data (Fig. 5a; Supplementary video S3) and quantified it with a Farazdaghi–Harris function [38], *y* = (*a* + *bx*^*c*^)^*−*1^, with fitted coefficients *a* = −20.727, *b* = 30.255, and *c* = −0.311. Among the candidate functions examined (SI Section IV), this profile narrows approximately linearly toward the outlet over much of the nasus length.

To determine how this geometry affects pressure-driven ejection, we performed computational fluid dynamics simulations of Herschel–Bulkley flow through the reconstructed nasus and through comparison nozzles with linear, quadratic, and other tapered profiles (Fig. 5b,c). All nozzle profiles had the same length and inlet and outlet diameters and were simulated using identical wall boundary conditions and material parameters; internal volume and taper ratio varied with profile shape. The reconstructed nasus and linear taper produced similar velocity fields and pressure losses, whereas the more strongly curved profiles generated larger pressure drops under the tested conditions.

**FIG. 5.**
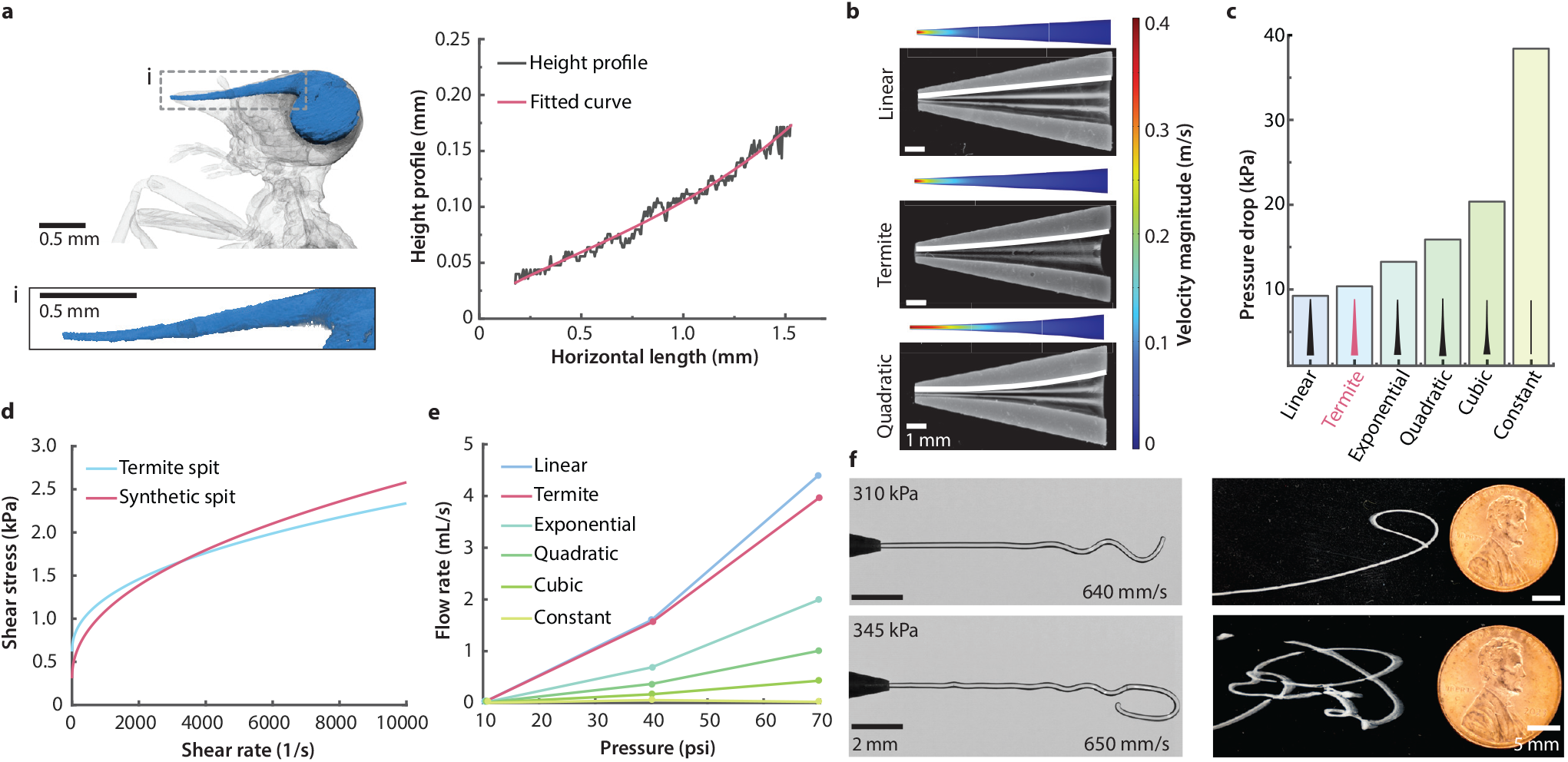
The near-linear nasus taper minimizes the pressure cost of ejecting a shear-thinning secretion. **a**. Internal nasus profile reconstructed from micro-CT data and compared with candidate fitted functions. **b**. Simulated velocity fields for Herschel–Bulkley flow through the reconstructed nasus and comparison nozzle profiles. Cross sections of the corresponding scaled, three-dimensionally printed nozzles are shown below. **c**. Simulated pressure drop through each nozzle profile for a Herschel–Bulkley flow index of *n* = 0.2. **d**. Herschel–Bulkley model of the natural secretion (parameters inferred from extensional rheometry and ejection kinematics; SI Section IV) compared with measured steady-shear rheology of the Pluronic-based synthetic analog. The two exhibit similar yield-stress and shear-thinning behavior over the shear-rate range relevant to extrusion. **e**. Measured flow rate of the synthetic analog through the fabricated nozzle profiles as a function of applied pressure. **f**. Representative horizontal ejection of the synthetic analog through a termite-shaped nozzle at different ejection conditions of applied pressure. The filament forms curved and looped deposits after contacting the substrate. Scale bars are 0.5 mm, 1 mm, 2 mm and 5 mm for the termite micro-CT reconstruction, fabricated nozzle cross sections, nozzle ejection, and synthetic secretion images, respectively.

The natural secretion and synthetic hydrogel were both described using a Herschel–Bulkley relation (Fig. 5d). For the synthetic hydrogel, the parameters were obtained by fitting measured shear stress against shear rate (Methods III). The relation 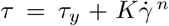, where *τ*_*y*_ is the yield stress, *K* is the consistency index, *n* is the flow index, and 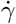 is the shear rate, allows comparison of their non-Newtonian behavior at extrusion shear rates.

Shear-thinning behavior corresponds to 0 *< n <* 1, and the resulting parameters give *n* = 0.46 and *n* = 0.44 for the termite secretion and synthetic hydrogel, respectively. Under pressure-driven flow, the high shear rate near the nozzle walls reduces the apparent viscosity, lowering the resistance to ejection relative to a Newtonian fluid of comparable low-shear viscosity. The synthetic analog reproduces this shear response but not the extensional hardening or long-term aging of the natural secretion.

We fabricated scaled versions of each nozzle profile and measured the flow rate of the synthetic analog under controlled applied pressures (Supplementary video S3). The experimental measurements reproduced the principal trend predicted by the simulations: the termite-shaped and linearly tapered nozzles produced comparable flow rates, whereas the curved profiles passed less flow at the same pressure (Fig. 5e). Agreement between the computational and physical models supports the conclusion that the near-linear nasus profile facilitates pressure-efficient extrusion of the shear-thinning secretion.

A termite-shaped nozzle enlarged by a factor of 20 also reproduced qualitative features of the natural ejection (Fig. 5f). When discharged horizontally above a substrate, the synthetic filament remained continuous in flight and formed curved or looped deposits upon contacting the surface. The detailed pattern varied across ejection conditions and trials, consistent with the sensitivity of filament deposition to small perturbations. Related tapering principles are used to reduce pressure losses during direct-ink writing of non-Newtonian materials [39], and supplementary proof-of-concept experiments extend the present system to extrusion of termite-inspired structures (Fig. S14).

### Nozzle height and orientation set lasso coverage in a scaled analog

Using the same shear-rheologically matched hydrogel analog and scaled termite-shaped nozzle, we constructed a mechanical ejection system to explore kinematic and geometric conditions beyond those accessible in living termites (Fig. 6a,b). The nozzle oscillated reciprocally while continuously ejecting the hydrogel, reproducing the two ingredients of the natural behavior, a continuous jet and a reciprocal sweep, while allowing each control parameter to be varied independently. These experiments were designed as a design-space study inspired by termite behavior, rather than as a dynamically identical model of the natural ejection. The main difference is gravity: the Froude number *u*^2^*/gh*, with *g* the gravitational acceleration and *h* the fall height, is of order 10 for the termite jet over the millimeter distances at which coiling was measured, but only 0.2–1.6 over the 10–100 mm heights of the scaled rig, so the scaled system explores a regime in which gravity competes with nozzle motion and coiling in shaping the deposit.

**FIG. 6.**
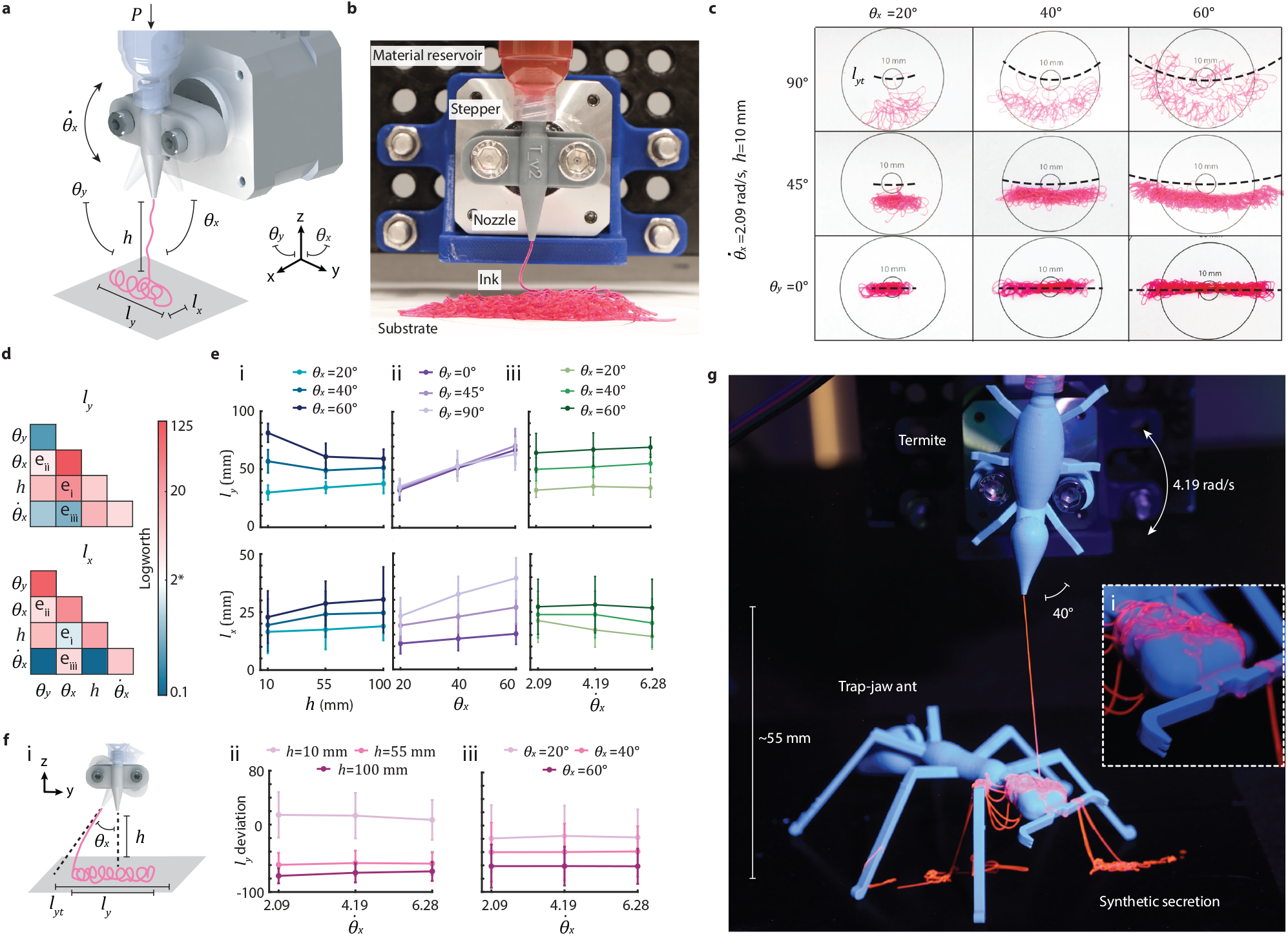
Nozzle height, orientation, and sweep jointly set synthetic lasso coverage. **a**. Schematic of the oscillatory-ejection system and the termite-shaped internal nozzle geometry. **b**. Experimental apparatus. The nozzle oscillates while extruding the synthetic secretion analog, and the resulting deposit is characterized by its width *l*_*y*_ and depth *l*_*x*_. **c**. Representative liquid-lasso deposits generated at different pitch angles *θ*_*y*_ and yaw angles *θ*_*x*_, with angular velocity 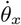 and nozzle height *h* held constant. **d**. Summary of the main and interaction effects of the four experimental factors on deposit dimensions. Statistical significance is represented by the LogWorth, *−* log_10_(*p*). **e**. Representative interaction effects between: **i**, nozzle height and yaw angle; **ii**, pitch angle and yaw angle; and **iii**, angular velocity and yaw angle. **f**. Percentage difference between the measured deposit width and the geometrically predicted sweep width as a function of angular velocity for different nozzle heights. **g**. Termite-shaped nozzle depositing a synthetic liquid lasso onto a model trap-jaw ant. Inset: looped and overlapping filaments deposited across the model head and mandibles. Error bars in e and f represent standard deviations.

We used a full-factorial design with four factors at three levels each: nozzle height above the substrate *h* = 10, 55, and 100 mm; pitch angle *θ*_*y*_ = 0, *π/*4, and *π/*2; yaw angle *θ*_*x*_ = *π/*9, 2*π/*9, and *π/*3; and angular velocity 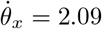, 4.19, and 6.28 rad/s. Here *θ*_*y*_ = 0 corresponds to ejection vertically downward onto the substrate and *θ*_*y*_ = *π/*2 to horizontal ejection, the orientation of the natural behavior (Fig. S11c).

The deposited structures varied in width, depth, and filament density. We quantified the planform width *l*_*y*_ and depth *l*_*x*_ as measures of the spatial footprint produced by each parameter combination (Fig. 6b,c). Both pitch angle and yaw angle affected the deposit dimensions. Increasing *θ*_*x*_ increased the nominal lateral sweep of the nozzle, while increasing *θ*_*y*_ redirected a larger component of the filament trajectory out of the plane of oscillation. Consequently, combinations of large pitch and yaw angles generally produced broader and deeper deposits.

For comparison with the measured width, we calculated the geometrically predicted sweep width, *l*_*y,t*_ = 2(*h* + *l*_*n*_) tan *θ*_*x*_, where *l*_*n*_ = 19.55 mm is the distance from the axis of rotation to the nozzle outlet. The difference between the measured width *l*_*y*_ and *l*_*y,t*_ separates the direct effect of nozzle motion from additional displacement caused by filament deformation, coiling, and gravitational deflection.

The factorial analysis identified significant main effects of nozzle height, pitch angle, and yaw angle, together with several interaction effects (Fig. 6d). Significant interactions included 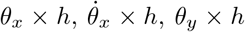, and *θ*_*y*_ × *θ*_*x*_, indicating that the influence of any single parameter depended on the configuration of the others. In particular, nozzle height altered the effects of both pitch and yaw on the deposit dimensions, while simultaneous increases in *θ*_*y*_ and *θ*_*x*_ produced larger two-dimensional footprints than either angular change alone (Fig. 6c,e).

Increasing the nozzle height *h* changed the partitioning of the deposit between width and depth. At larger *h*, deposits tended to converge in width but diverge more strongly in depth across angular configurations (Fig. 6e). This behavior is consistent with the longer unsupported filament length increasing the opportunity for gravitational deflection and out-of-plane motion before substrate contact. By contrast, extrusion close to the substrate promoted local coiling near the contact point, extending the measured deposit beyond the nominal geometric sweep (Fig. 6f).

To test whether the excess deposit width arose primarily from inertial lag during nozzle oscillation, we calculated the percentage excess Δ*l*_*y*_ = 100 (*l*_*y*_ −*l*_*y,t*_)*/l*_*y,t*_. The excess width did not increase systematically with angular velocity over the tested range (Fig. 6f). Inertial lag of the filament behind the accelerating nozzle therefore cannot explain the excess; its dependence on height and orientation, together with the coiling observed in the termite (Fig. 3), points to filament coiling and deformation near the substrate.

As a qualitative demonstration of the resulting coverage, the oscillating system deposited overlapping loops across the head, mandibles, and surrounding region of a model trap-jaw ant (Fig. 6g; Supplementary video S3). The deposited footprint extended both within and outside the plane swept directly by the nozzle, illustrating how reciprocal motion and filament instability can distribute material over a target larger than the ejecting organism.

## DISCUSSION

The liquid-lasso defense of *Nasutitermes* termites integrates morphology, kinematics, and time-dependent material behavior to overcome a predator–prey size asymmetry (Fig. 1 and Supplementary video S1). Rather than relying on a single optimized feature, the system operates sequentially: a shear-thinning secretion is rapidly ejected through a tapered nasus, lateral head oscillation couples with rope-coiling instability to expand the deposition footprint, and strain hardening followed by aging stabilizes the resulting loops and bridges. Defensive effectiveness therefore emerges from coordination among processes acting at different stages of deployment.

The secretion complements this spatial amplification through a time-evolving mechanical response. Shear thinning lowers the apparent viscosity during flow through the narrow nasus, whereas strain hardening under extension after deposition raises the resistance to filament breakup. Subsequent aging further stabilizes the material and ultimately produces a solid-like adhesive This progression allows the secretion to satisfy competing functional requirements: it flows during ejection, remains filamentary during deposition, resists removal during predator movement and grooming, and hardens after contact. Monoterpene loss may contribute to this solidification process [12], although the chemical mechanisms governing this process remain to be determined.

Downstream of the nasus, the termite exploits (rather than suppresses) fluid instability. Soldiers actively scan the nasus laterally, while the deposited filament passively buckles into coils, meanders, and overlapping strands. Together these motions make the termite a reciprocal liquid sewing machine [28, 29], in which muscular motion defines the region traversed by the jet and filament instability redistributes material within and beyond it. The scaled experiments further show that coverage depends on interactions among nozzle height, orientation, angular excursion, and oscillation rate. The natural behavior thus appears to occupy a coupled geometric and kinematic regime that generates broad, irregular coverage.

These material and kinematic mechanisms act on the ant predators in complementary ways. Coiling increases the probability of contacting multiple body regions, strain hardening keeps those contacts intact while the ant struggles (its own pulling may stiffen them further), and solidification converts them into persistent mechanical constraints. The behavioral signature is a large increase in grooming that fails to clean the deposit (SI Section II). Whether contamination of the antennae, the principal chemosensory organ of ants [40], and of the eyes also degrades chemosensory or visual performance was not directly assessed here and remains an open question.

The nasus contributes primarily to the ejection stage. Its internal profile approximates a near-linear taper, and both simulations and scaled experiments showed pressure losses comparable to those of a linear nozzle and lower than those of the other curved profiles tested. Combined with the secretion’s yield-stress and shear-thinning behavior, this geometry keeps the pressure cost of ejection low. Whether the nasus is a global geometric optimum, and what pressure the gland supplies, remain untested. Nonetheless, the current data indicate that the biological nozzle occupies a broad, low-resistance region of the tested design space.

The rheological differences observed between *N. corniger* and the unidentified *Nasutitermes* sp. suggest that secretion mechanics may vary among species. Stronger strain hardening and arrested breakup in *N. corniger* could reflect differences in secretion chemistry, diet, colony condition, nesting ecology, or predator communities. Broader comparative sampling under standardized conditions will be needed before such differences can be interpreted as species-specific adaptations.

By coupling active scanning with passive instability and delayed solidification, the termite converts a nanoliter-scale secretion into a body-scale adhesive network. These principles may inform engineered systems requiring low-pressure extrusion, broad deposition, and time-programmed solidification of non-Newtonian materials, including oscillatory direct-ink writing [41, 42], deployable adhesive networks, targeted delivery, and soft robotic fluid deployment. Supplementary printing experiments (Fig. S14) provide an initial proof of concept for extending these ideas toward fabrication of termite-inspired structures. But the broader design principle is the coordinated use of nozzle geometry, imposed motion, and evolving rheology to control material deployment across multiple stages.

## METHODS

### I. Field extensional rheology

To characterize the extensional response of the natural secretion, we mechanically induced multiple soldier termites to deposit secretion onto a circular metal plate. A second plate was lowered from above to contact the secretion and form a liquid bridge with an initial plate separation *h*_0_ *<* 0.5 mm. The bridge was allowed to relax for up to 2 min before the upper plate was raised to a final separation *h*_*f*_, imposing uniaxial extension. The subsequent capillary thinning was recorded using high-speed imaging.

The filament diameter *D*(*t*) was measured at the axial midplane, *z* = *h*_*f*_ */*2, and normalized by the diameter *D*_1_ immediately after extension. Preliminary experiments were used to identify an extension range that generated capillary thinning without either prolonged under-extension or immediate failure caused by excessive extension.

During the early viscocapillary regime, the apparent extensional viscosity *η*_*E*,0_ was estimated by fitting Eq. (1) over *t <* 2 s with *γ* = 30 mN/m (SI Section III). During the subsequent elastocapillary regime, the relaxation time *λ* was obtained by fitting Eq. (2).

Since the zero-shear viscosity of the natural secretion could not be measured independently, the apparent extensional viscosity was normalized by its early-time value, 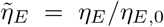. Hencky strain, extension rate, and Deborah number were calculated as *ε* = 2 ln(*D*_1_*/D*), 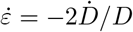, and 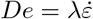.

To characterize aging, capillary-thinning measurements were repeated on deposited secretion after different elapsed times. Since the secretion has a high surface-to-volume ratio and may evolve through evaporation or other physicochemical processes, the aging time was recorded from initial deposition.

### II. Preparation of the synthetic secretion analog

A Pluronic F-127 hydrogel was prepared by gradually adding Pluronic F-127 (Sigma–Aldrich) to chilled water to obtain a 30:70 polymer-to-water mass ratio. The mixture was stirred using a magnetic stirrer (SH-2, Anpooz) until homogeneous.

For oscillatory-lasso experiments, a red water-based fluorescent dye (Risk Reactor) was added at a dye-to-hydrogel mass ratio of 1:50. The mixture was homogenized in a speed mixer (FlackTek) at 2500 rpm for 2 min.

### III. Steady-shear characterization of the synthetic analog

Steady-shear measurements were performed using an Anton Paar MCR 302 rheometer. The unfilled hydrogel was tested using a 25-mm parallel-plate geometry. Soil-containing formulations used in the supplementary printing experiments were tested using an 8-mm sand-blasted parallel-plate geometry. Shear-rate sweeps were performed from 0.1 to 10^4^ s^*−*1^, and the corresponding shear stress and apparent viscosity were recorded.

The shear-stress data were fitted using the Herschel– Bulkley relation 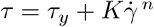 defined in Results.

### IV. Fabricated-nozzle flow measurements

Scaled nozzle profiles were fabricated for experimental comparison with the computational models. All profiles had the same nozzle length and inlet and outlet diameters; internal volume and taper ratio varied according to the profile shape.

The flow rate of the synthetic analog through each nozzle was measured at applied pressures of 10, 40, and 70 psi. Material was collected for 10, 1, and 0.5 s, respectively, to maintain measurable output volumes across the pressure range. Up to five replicate measurements were performed for each nozzle and pressure condition.

For the oscillatory-lasso experiments, the applied pressure was varied until the measured filament velocity was 391 mm*/*s, within the range measured for natural termite ejection. This condition was obtained at an applied pressure of 17.5 psi.

### V. Representative jetting through the termite-shaped nozzle

Representative ejection behavior was recorded using a termite-shaped nozzle enlarged by a factor of 20, with an outlet diameter of 0.64 mm. The synthetic analog was ejected at applied pressures of 45 and 50 psi (310 and 345 kPa). A pneumatic solenoid valve was opened for 0.1 s to generate each pulse. Ejection was recorded using a FASTCAM SA5 high-speed camera (Photron) at 18,600 frames/s.

### VI. Scaled oscillatory-lasso experiments

The scaled oscillatory-ejection apparatus consisted of a stepper motor (17HS4401S) mounted on a motion gantry (Aerotech). Pressure was supplied using an Ultimus V pressure controller (Nordson EFD). The motor was driven by a DM542T motor driver (StepperOnline) controlled by an Arduino Uno. A pneumatic solenoid valve (VQD-1151, SMC) synchronized the onset of material extrusion with the start of nozzle motion.

The 20× scaled termite-shaped nozzle oscillated reciprocally between positive and negative yaw angles for three cycles before the motor and pneumatic valve were stopped simultaneously. The experimental factors were nozzle height above the substrate *h*, nozzle pitch angle *θ*_*y*_, yaw angle *θ*_*x*_, and angular velocity 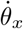. A full-factorial design was used with *h* = 10, 55, and 100 mm; *θ*_*y*_ = 0, *π/*4, and *π/*2; *θ*_*x*_ = *π/*9, 2*π/*9, and *π/*3; and 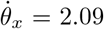, 4.19, and 6.28 rad/s.

### VII. Image analysis and statistics for oscillatory-lasso experiments

Deposits from the oscillatory-lasso experiments were imaged and analyzed using ImageJ. The planform width *l*_*y*_ and depth *l*_*x*_ were measured separately. The geometrically predicted sweep width *l*_*y,t*_ and the percentage difference Δ*l*_*y*_ were calculated as defined in Results.

Factorial analyses were performed in JMP. Since both width and depth exhibited heteroscedasticity, and the depth measurements also deviated from normality, Box– Cox transformations were applied before analysis. The transformation parameters were *λ* = 0.505 for width and *λ* = 0.262 for depth. Main effects and interaction effects of *h, θ*_*y*_, *θ*_*x*_, and 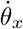 were then evaluated using the full-factorial model.

### VIII. Termite-inspired printing and compression testing

The following procedures support the supplementary printing demonstration. Tropical soil (Organic Mechanics) was sieved and mixed with the Pluronic F-127 hydrogel at soil-to-hydrogel mass ratios of 1:1, 1:5, and 1:10. Samples were mixed in a speed mixer at 2500 rpm for 2 min and centrifuged in 10- or 30-mL syringes at 5000 rpm for 1 min using a ProcessMate 5000 centrifuge (Nordson EFD). Compression-test specimens were printed using a termite-shaped nozzle enlarged by a factor of 50. The nominal specimen dimensions were approximately 15 mm in diameter and 11 mm in height. The 1:1, 1:5, and 1:10 soil-to-hydrogel formulations were printed at 26, 14, and 11.3 psi, respectively, using a feed rate of 10 mm/s. After printing, specimens were either dried under ambient conditions or heated overnight at 40, 70, or 100^*°*^C, depending on the experimental condition. Compression tests were performed using a universal testing machine (MTS) at a displacement rate of 0.1 mm/s. Force and displacement were recorded and converted to engineering stress and strain.

## DATA AVAILABILITY

All data needed to evaluate the conclusions in the paper are present in the paper and/or the Supplementary Materials. High-speed videos, rheology data, and analysis scripts are available at https://github.com/echallita/Termites_defense. Micro-CT reconstructions, simulation files, and the analysis scripts will be deposited in Zenodo with a DOI before publication.

## CODE AVAILABILITY

Analysis scripts are included in the repository above.

## ACKNOWLEDGEMENTS

S.B. acknowledges funding support from Schmidt Sciences LLC. C.M. acknowledges funding support from the Georgia Tech - University Center of Exemplary Mentoring (GT-UCEM) Fellowship program and Georgia Tech’s President’s Fellowship. P.R. acknowledges an Eckert Postdoctoral Fellowship from the Georgia Tech Foundation. J.M. gratefully acknowledges support from the Jean-Jacques and Felicia Lopez-Loreta Foundation through the 2020 Prize.

## AUTHOR CONTRIBUTIONS

EJC: Conceptualization, Data curation, Formal analysis, Funding acquisition, Investigation, Methodology, Project administration, Resources, Software, Supervision, Validation, Visualization, Writing - Original Draft, Writing - Review & Editing; DCA: Conceptualization, Data curation, Formal Analysis, Investigation, Methodology, Validation, Visualization, Writing – original draft; PR: Data curation, Investigation, Methodology, Validation, Writing – Reviewing and editing; JM: Conceptualization, Funding acquisition, Methodology, Project administration, Supervision, Visualization, Writing – original draft; CM: Visualization, Writing – Reviewing and editing; SB: Conceptualization, Funding acquisition, Investigation, Project administration, Resources, Supervision, Validation, Writing - Original Draft, Writing - Review & Editing; JSH: Investigation, Methodology, Writing - Review & Editing; PS: Investigation, Writing - Review & Editing; DAN: Resources, Investigation, Writing - Review & Editing.

## COMPETING INTERESTS

The authors declare no competing interests.

## Notes

https://github.com/echallita/Termites_defense

### Competing Interest Statement

The authors have declared no competing interest.

