## Supplementary Information for "Viscoelastic liquid lassos extend the defensive reach of termite soldiers"

### Contents

I. **Data collection and analysis** 1
- A. Geographical location and field setup 1
- B. Kinematics of ejection and head movement 2
- C. SEM of termites and ants 2

II. **Termite–ant interactions** 2
- A. Field assays of ant grooming after termite exposure 2
- B. Statistical analysis of ant behavior 2
- C. Grooming response to termite exposure 2

III. **Fluid properties** 3
- A. Surface tension and density of the secretion 3
  - 1. Surface tension ($\gamma \approx$ 25–35 mN/m) 3
  - 2. Density ($\rho \approx$ 850–950 kg/m$^3$) 3
- B. Liquid rope coiling 3
- C. Capillary-breakup extensional rheology of termite secretion 3

IV. **Nasus geometry, flow models, and synthetic analog** 4
- A. Termite nozzle shape 4
- B. Analytical flow model 4
- C. Numerical flow model 5
- D. Shape functions 6
- E. Parametric studies 6
  - 1. Effect of material properties 6
- F. Flow regime in the natural and scaled nozzles 7
- G. Rheology of the synthetic secretion analog 8
  - 1. Ejection of the synthetic analog through the scaled nozzle 8
  - 2. Extrusion printing with the termite-shaped nozzle 8

V. **Supplementary videos** 8

VI. **Supplementary figures** 9

VII. **References** 22

**Supplementary References** 22

### I. Data collection and analysis

#### A. Geographical location and field setup

Fieldwork was conducted at Los Amigos Research Station and Finca Las Piedras Research Station in Madre de Dios, Peru; La Selva Biological Station in Costa Rica; Barro Colorado Island in Panama; and at Osa Conservation on the Osa Peninsula, Costa Rica. The picture of the termite interacting with the trap jaw ant was taken in Fond Doux, Saint Lucia (13.844, -61.036).Termites were identified by their distinctive nasus and located by systematically scanning arboreal nests and associated canal networks on trees.

*Nasutitermes* sp. were sampled at Finca Las Piedras Research Station and Los Amigos Biological Station in the Madre de Dios region of Peru, while *Nasutitermes corniger* (Motschulsky, 1855) [1] specimens were collected from Barro Colorado Island, Panama. All studied species were arboreal and characterized by long, external canals extending from their nests to lower regions of the tree trunk, facilitating access to foraging and defensive sites.

Following termite localization, we deployed a portable high-speed imaging setup powered by a generator. High-speed videography was primarily conducted using a Photron Mini AX camera equipped with a Canon MP-E 65 mm macro lens at 1×–3× magnification. Additional datasets were collected using a FASTEC V5 high-speed camera for black-and-white imaging. Illumination was provided by a field-portable ZAILA high-intensity light system powered by portable battery packs to ensure sufficient lighting for high-speed capture.

After observation and videography, termite specimens were collected and preserved in 200-proof ethanol for species identification and further analysis.

#### B. Kinematics of ejection and head movement

For the jet ejection experiment, we tracked the jet tips as they exited the termite nozzle and before hitting a substrate. We examined two scenarios: 1) termites in their natural setting, mechanically disturbed by external substrates (e.g., glass slides, leaves, lens caps, knives, tweezers), and 2) termites held with tweezers while their antennae were carefully stimulated against a glass slide. We observed that termites can eject multiple times similar to *Nasutitermes exitiosus* (Hill, 1925) [2] reported by Eisner. In the tweezers-held scenario, termites typically eject initially when picked up from the tree. Consequently, the filmed ejections are often secondary or tertiary. Despite this, we noted no significant difference in jet speeds between initial and subsequent ejections.

Jet exit speeds were recorded at 10,000 fps, while head movements were captured at 5,000 fps and analyzed at 1,000 fps (downsampled). We applied a moving average filter (size 3 for ejection data, size 5 for head movements) to smooth the raw data. Analysis was performed using MATLAB and ImageJ (FIJI).

We measured the 2D location of the jet tip $P(x, y)$ before substrate impact. Displacement was calculated using the Euclidean distance formula $d = \sqrt{(x_f - x_e)^2 + (y_f - y_e)^2}$. Ejection speed ($V_d$) was derived by averaging the speed measurements: $V_d = \partial_t\, d$.

For head movements, we tracked the nozzle tip post-stimulation. We consistently observed the initial movement of the head of the termites toward the stimulated antennae. Although termite heads move in 3D (confirmed by side-view videos), our analysis focused on instances where the head remained in focus and therefore the data presented in the paper corresponds to the movement of the heads where it stays mostly in a 2D plane. This approach introduced an error range of $88 - 150\mu m$, based on the depth of field of the Canon MPE 65 mm lens at 2×-3× magnification. The angular head movement ($\theta$) was calculated relative to the axis running along the termite's head.

#### C. SEM of termites and ants

Termite and ant specimens were euthanized via 80% ethanol and stored in vials before bringing back from the field. Before imaging and to preserve their natural morphology, specimens underwent ambient dehydration during storage and were not subject to chemical dehydration or sputter coating with a conductive film. During imaging, samples were mounted on carbon conductive tabs to facilitate electron dissipation. Images were taken at 5kV using a Phenom Pro Desktop SEM (Nanoscience Instruments, Arizona, United States).

### II. Termite–ant interactions

#### A. Field assays of ant grooming after termite exposure

To quantify the behavioral response of ants to termite exposure, we conducted three-stage field assays with three sympatric ant taxa observed on or near *Nasutitermes* nests and known to include opportunistic termite predators: the army ant *Eciton harpax*, *Camponotus* sp., and *Crematogaster* sp. [3, 4]. Assays were run in plastic arenas measuring 5 × 5 × 1 in. Each ant was first observed alone in a clean arena for 1 min (Pre), transferred for 1 min to an encounter arena containing approximately 20–30 soldier termites, 3–5 worker termites, and fragments of termite nest material, and then transferred to a third clean arena for 1 min (Post). Control ants underwent the same three-stage protocol without termites (Fig. S2a). Arenas were sanitized with 70% ethanol between trials.

We tested 10 ants in the experimental group and three control ants for each ant taxon. Ant behavior was annotated from video recordings using BORIS. We recorded the onset and termination of self-grooming events involving the antennae or body during the Pre and Post observation periods. For each ant, the *cleaning ratio* was calculated as the total time spent grooming divided by the total observation time, and the *bout frequency* as the number of discrete grooming events during the observation period.

#### B. Statistical analysis of ant behavior

Cleaning ratio and grooming-bout frequency were analyzed using linear mixed-effects models. State (Pre or Post), ant taxon (*Eciton harpax*, *Camponotus* sp., or *Crematogaster* sp.), Group (Control or Experimental), and their interactions were included as fixed effects, with ant identity included as a random intercept to account for repeated Pre and Post observations of the same individual.

Post-hoc comparisons between Pre and Post states were performed within each Taxon×Group combination using estimated marginal means. The resulting $p$-values were adjusted for multiple comparisons using the Benjamini–Hochberg procedure. Analyses were conducted in R using the `lme4`, `lmerTest`, and `emmeans` packages.

#### C. Grooming response to termite exposure

We compared exposed and control ants using linear mixed-effects models. Both cleaning ratio and bout frequency exhibited significant State×Group interactions (cleaning ratio: $F_{1,32.8} = 7.12$, $p = 0.012$; bout frequency: $F_{1,33} = 7.98$, $p = 0.008$), indicating that the

Pre-to-Post change differed between termite-exposed and control ants. Within the control group, no taxon showed a significant Pre-to-Post change in either measure (all post-hoc contrasts $p > 0.59$), indicating that handling and repeated observation alone did not measurably alter grooming.

Among termite-exposed ants, *Camponotus* sp. increased their mean cleaning ratio from 4.0% before the encounter to 28.6% afterward, an increase of 24.6 percentage points ($p = 0.002$). *Crematogaster* sp. increased their cleaning ratio from 0.7% to 35.0%, an increase of 34.3 percentage points ($p < 0.001$) (Fig. S2b). *Eciton harpax* showed no significant change in cleaning ratio, increasing from 4.1% to 9.0% ($p = 0.512$), but its mean bout frequency increased from 1.4 to 4.0 bouts ($p < 0.001$). Thus, *E. harpax* responded through shorter, more frequent grooming events rather than prolonged grooming sessions. Grooming-bout frequency increased after termite exposure in all three taxa (Fig. S2c).

The observed grooming was concentrated on the antennae, head, and limbs, including structures central to chemosensation and grooming [5]. Scanning electron micrographs provided direct evidence that the secretion reached these regions. Following termite interactions, *Neoponera apicalis* (Latreille, 1802) [6] and *Odontomachus bauri* Emery, 1892 [7] exhibited secretion on the eyes, antennae, cuticular surfaces, and grooming setae and brushes (main text Fig. 2c,d). The secretion frequently remained attached to fine surface structures, and grooming appeared in some observations to redistribute the material onto the ants' own cleaning apparatus rather than remove it completely. These behavioral and microscopic observations are consistent with persistent contamination of sensory and grooming structures, although direct impairment of sensory performance was not measured.

The secretion also imposed mechanical constraints. Eisner et al. reported that arthropods exposed to *N. exitiosus* secretion could become physically immobilized within minutes [8]. We observed comparable effects in the present system. Small ants, including *Crematogaster* sp., moved with difficulty after termite encounters, while larger trap-jaw ants were occasionally tethered to the substrate by filamentous strands of secretion (Supplementary video S1). In overnight observations, some ants remained alive but entangled in material that had spread across the body and hardened into a rigid coating. These observations are consistent with the rheological progression measured in Section III: freshly deposited filaments resist extension and breakup, while subsequent aging converts temporary bridges into persistent solid-like restraints.

### III. Fluid properties

#### A. Surface tension and density of the secretion

##### 1. Surface tension ($\gamma \approx 25$–$35\ mN/m$)

Termite secretion is primarily composed of $\alpha-$pinene and limonene having surface tensions of 25.87 $mN/m$ [9] and 27 $mN/m$ [10] at $25°C$. Assuming that the surface tension is modulated primarily by these two solvents, we would expect a surface tension of around 25 to 35 $mN/m$. This hypothetical range and the actual value could differ based on precise composition, temperature, chemical composition and purity of the fluid itself.

##### 2. Density ($\rho \approx 850$–$950\ kg/m^3$)

The termite secretion may be described as an organic terpenoid-resin mixture. Its density may be inferred from dominant components: limonene ($841 kg/m^3$), $\alpha$-pinene ($858 kg/m^3$), and $\beta$-pinene ($872\ kg/m^3$).

#### B. Liquid rope coiling

Termites were carefully held with tweezers and gradually approached to glass slides. To induce fluid ejection, we gently pressed their heads against the glass, ensuring that their antennae were mechanically stimulated. The ejection process was recorded at 10,000 frames per second (fps) using 2x and 3x magnification with a Canon MPE 65 mm lens, providing a temporal resolution of $dt = 1/10,000\ s$.

We measured the distance between the nozzle ($H$) and the substrate, and corrected for this distance as the fluidic coil accumulated. Additionally, we measured the speed of ejection before the fluid hit the surface, though measuring the speed of the jet after impact proved challenging given the continuous nature of the jet. The fluctuations observed in the coiling frequency be attributed to variations in ejection speed and the termites' heads moving in and out of the camera's focus. However, we only analyzed the coils where the fluid remained within in focus the frame and within a 20° angle from the vertical.

#### C. Capillary-breakup extensional rheology of termite secretion

Capillary-breakup extensional rheology experiments were conducted in the Amazon rainforest and the Barro Colorado Island using a custom-built setup inspired by CaBER techniques [11]. The apparatus consisted of a stepper-motor-driven linear translation stage; additional experiments were performed using a manually actuated micro–linear stage.

For experiments conducted in the Amazon, the fluid was confined between two parallel stainless-steel hex key surfaces. Experiments performed at Barro Colorado Island (BCI) used two parallel glass slides as substrates. Termites were collected by gently positioning an individual near the lower substrate and inducing secretion. Immediately after secretion, the upper substrate was slowly brought into contact with the fluid to form a thin fluid bridge, which was allowed to equilibrate for approximately 10 s before extension.

The initial gap height was $h_0$ with an initial filament diameter $D_0 \sim 1$ mm. The filament was subsequently stretched to a final gap height $h_f$, corresponding to initial and final aspect ratios $\Lambda_0 = h_0/D_0$ and $\Lambda_f = h_f/D_0$, respectively. The initial gap height was chosen such that $h_0$ remained smaller than the capillary length, $l_c = \sqrt{\gamma/(\rho g)}$. Due to the limited fluid volume, the final filament diameter varied slightly across trials.

Filament thinning and breakup dynamics were recorded at 500 fps using a Photron high-speed camera equipped with a Canon MP-E 65 mm macro lens, yielding a spatial resolution of approximately 9 $\mu$m. Filament diameter was measured at the mid-plane of the filament, $h_{1/2} = h_f/2$. Only experiments exhibiting symmetric filament breakup were included in the analysis. To minimize drying or curing effects, only trials with capillary breakup times in the range $4 < t_c < 14$ s were analyzed.

Ambient temperature and relative humidity were monitored throughout experiments and ranged from 30°C to 34°C and 80–90%, respectively. Each experimental configuration required approximately 2–3 minutes of preparation and data collection.

### IV. Nasus geometry, flow models, and synthetic analog

#### A. Termite nozzle shape

We analyzed the termite head shape with a focus on the nozzle. The microCT images were prepared as follows for analysis via Python code (Figure S5a,b). In Step 1, the original images from the microscopes were prepared using image processing software to ensure only the respective regions of interest, i.e., the inner nozzle volume, was highlighted in color. The nozzles were aligned horizontally with the termite head on the right and imported in the Python code. In Step 2, the images were manually cropped to focus on only the nozzle; all successive steps were executed autonomously by the code without the need for human intervention. A part of the fluid container behind the nozzle was kept to visualize the transition. In Step 3, we isolated a predetermined Hue Saturation Value (HSV) color range with lower and upper bounds of [100, 150, 50] and [140, 255, 255], respectively. In Step 4, these images were converted to grayscale for later height extraction. In Step 5, a grayscale threshold value of 80/255 was defined to increase contrast and provide clearer results. In Step 6, the images were sliced into columns of one pixel width. In Step 7, all pixels above the threshold were counted to determine the nozzle height as a function of the nozzle length. The actual height was calculated with a scaling factor determined for each image individually by dividing the number of pixels along the scale bar by the length of the scale bar in millimeters. The full details, Python code, and additional parameter values used can be found in the SI.

Due to microscopic and numerical inaccuracies, e.g., a stair-stepping effect (Fig. S5c,d), and for the following parametric analysis, curve fitting was conducted on the experimental nozzle height data. As we are not aware of a specific function representing the underlying geometry or physics of the termite nozzle shape, first, a general qualitative fit with over 180 diverse fitting functions taken from the OriginLab database was evaluated.[12] Five best-fitting functions were selected to be investigated quantitatively (Table S1). Two power law models provided the lowest root mean square errors (RMSE): The Farazdaghi and Harris Model,[Ratkowsky (1989, page 89) and Seber (1989, page 362)] typically used in yield-density studies, and the LangmuirEXT2 Model, which has been adapted from the Langmuir adsorption model. [13] For numerical simplicity, we selected the Farazdaghi and Harris Model.

#### B. Analytical flow model

To estimate the effect of geometry on various flow metrics, we developed a simplified analytical-numerical model for non-Newtonian liquids flowing through a nozzle of length $L$ with an arbitrary shape, defined by shape functions, between an inlet diameter $D_{\text{inlet}}$ and an outlet diameter $D_{\text{outlet}}$.

The liquid was modeled using the Herschel-Bulkley model, which adds a yield stress $\tau_y$ to the power law model, allowing input of the rheological parameters extracted from termite secretion and expansion towards common non-Newtonian materials for further evaluation, e.g., Bingham plastics in 3D printing. The Herschel-Bulkley model describes the relationship between shear stress $\tau$ and shear rate $\dot{\gamma}$ for a non-Newtonian fluid. It includes the yield stress $\tau_y$, below which the fluid does not flow. The equation is:

$$\tau = \tau_y + K \cdot \dot{\gamma}^n$$

where $K$ is the consistency coefficient and n is the flow behavior index.

The average velocity $v$ of the fluid at any cross-section of the nozzle is determined by the volumetric flow rate $Q$ and the cross-sectional area $A$ at that point:

$$v = \frac{Q}{A}$$

The cross-sectional area for a circular nozzle is calculated

TABLE S1. **Comparison of candidate functions fitted to the extracted termite nasus profiles.** RMSE values are reported for the two reconstructed nozzle profiles and as the mean across both profiles.

| **Model** | **Function,** $y =$ | **Head 1 RMSE** | **Head 2 RMSE** | **Mean RMSE** |
|---|---|---|---|---|
| Exponential: Exp3P2 | $e^{a+bx+cx^2}$ | 0.005 55 | 0.007 28 | 0.006 41 |
| Polynomial: Parabola | $a + bx + cx^2$ | 0.005 52 | 0.007 87 | 0.006 70 |
| Power: Farazdaghi–Harris | $(a + bx^c)^{-1}$ | 0.005 63 | 0.006 68 | 0.006 15 |
| Power: LangmuirEXT2 | $\left(a + bx^{c-1}\right)^{-1}$ | 0.005 63 | 0.006 68 | 0.006 15 |
| Power: Pow2P2 | $a(1 + x)^b$ | 0.005 52 | 0.008 37 | 0.006 95 |

using the diameter $D$ at that point:

$$A = \frac{\pi D^2}{4}$$

The shear rate at the wall $\dot{\gamma}_w$ is estimated based on the average velocity and the diameter of the nozzle. This is a simplified assumption and the actual profile can be more complex:

$$\dot{\gamma}_w = \frac{v}{\frac{D}{2}}$$

The pressure drop $\Delta P$ is calculated by integrating the wall shear stress over the length of the nozzle. The wall shear stress $\tau_w$ is related to the shear stress by the factor $\frac{4}{D}$ due to the perimeter of the cross-section for a circular duct, giving:

$$\Delta P = \int_0^L \frac{4\tau_w}{D} dx$$

Where $D$ is the diameter at any point $x$ along the nozzle and $\tau_w$ is the wall shear stress, calculated using the Herschel-Bulkley model and the estimated shear rate at the wall.

These equations are implemented and integrated numerically in a Python code to estimate the pressure drop. The actual behavior of a non-Newtonian fluid can be more complex, and for precise calculations, more detailed analysis or empirical data may be necessary. The shear rate approximation, in particular, is a simplification and might not accurately represent the actual flow profile near the wall. The analytical model was flow-rate controlled: a prescribed volumetric flow rate $Q$ was imposed, and the pressure drop required to sustain that flow was calculated for each nozzle geometry.

### C. Numerical flow model

The analytical model allows efficient calculation of core parameters, such as the pressure drop in the nozzle for a given flow rate, which can be used for further analytical studies, quick iterations of design parameters, and, to some extent, validation of theories, microscopic measurements, experiments, and numerical models, a comprehensive numerical model is required to calculate more complex variables with high dependencies on the complex fluids.

In the flow-rate-controlled analytical model the outlet velocity $v = Q/A$ is the same for every nozzle shape; only the pressure required to sustain it differs. In the termite, however, ejection is more likely pressure-driven, i.e., the gland sets the inlet pressure rather than the flow rate. The flow rate then becomes a dependent variable set by the pressure difference, the nozzle geometry, and the rheology. For Newtonian fluids in simple geometries the Hagen–Poiseuille relation would suffice; for a Herschel–Bulkley fluid in a tapered geometry the governing (Navier–Stokes) equations must be solved numerically.

The CFD module of COMSOL Multiphysics 6.1 was used to model steady, single-phase, incompressible flow through the nozzle geometries. The Navier–Stokes equations and conservation of mass were solved using a manually defined Herschel–Bulkley constitutive relation. The geometries were implemented as parametrized two-dimensional axisymmetric domains. All comparison nozzles, where applicable, had the same length and inlet and outlet diameters, while their internal volume and local taper varied according to the prescribed shape function. A physics-based fine mesh was used, with increased resolution near the walls and other regions of large velocity gradients.

Two related numerical protocols were used. Pressure-controlled simulations prescribed an inlet pressure and fixed the outlet pressure at zero gauge pressure to examine the resulting velocity and flow fields. For direct comparison of hydraulic resistance among nozzle geometries, the inlet pressure was varied until a common target flow rate, and therefore a common mean outlet velocity, was obtained; the pressure drop required to achieve this flow condition was then compared among geometries. This latter protocol underlies the pressure-drop comparison reported for the nozzle-shape study.

Owing to the strong nonlinearity of the Herschel–Bulkley constitutive relation, auxiliary continuation sweeps were used to improve numerical convergence. Depending on the parameter study, either the inlet pressure was ramped from a lower value to the target pressure or the flow-behavior index $n$ was ramped from $n = 1$ toward its target value.

### D. Shape functions

We selected nozzle 1 (termite head 1, Fig. S5a,c) as the reference geometry for the following study. The fitted Farazdaghi–Harris function is

$$y = (a + b \cdot x^c)^{-1}$$

with fitting coefficients $a = -20.727$, $b = 30.255$, and $c = -0.311$. Scaled to the actual nozzle dimensions, the profile has length $L = 1.355$ mm and inlet and outlet diameters $D_{\text{inlet}} = 0.173$ mm and $D_{\text{outlet}} = 0.032$ mm. For comparison with common nozzle shapes, we added the shape functions in Table S2.

TABLE S2. **Shape functions used to represent the termite nasus and comparison nozzle geometries.**

| Shape | Shape function |
|---|---|
| Termite nasus | $y = (a + bx^c)^{-1}$ |
| Constant diameter | $y = a$ |
| Linear | $y = a + bx$ |
| Quadratic | $y = a + bx + cx^2$ |
| Cubic | $y = a + bx + cx^2 + dx^3$ |
| Exponential | $y = ae^{bx}$ |

To simplify the analytical and numerical models, we shifted and oriented each profile so that the inlet lies at $x = 0$ and the outlet in the positive $x$ direction. We added an $x_{\text{offset}}$ parameter and a mirroring option about $x_{\text{max}}$, illustrated for the termite nozzle function below and in Fig. S6:

Original curve: $y = (a + b\,x^c)^{-1}$

Curve with $x$ offset: $y = (a + b\,(x + x_{\text{offset}})^c)^{-1}$

Mirrored curve: $y = (a + b\,(x_{\text{max}} - x + x_{\text{offset}})^c)^{-1}$

Nozzles with varying shapes are tailored for specific applications based on their unique fluid dynamic properties.

### E. Parametric studies

Using these shape functions, we compared several nozzle geometries under common geometric and flow constraints (SI Table S2). The Herschel–Bulkley parameters used to represent the secretion were inferred from the extensional-rheology measurements, the measured ejection kinematics, and pressure-constrained flow modeling (see below). The resulting reference parameter set was

$$\tau_y = 298 \text{ Pa}, \qquad k = 33 \text{ Pa s}^n, \qquad n = 0.46.$$

The flow rate was calculated using the jetting diameter of $D_j \sim 25\mu m$, and the velocity at the outlet $v = 400mm/s$ from the experiments to be $Q = 1.96 \cdot 10^{-10} m^3/s$.

#### 1. Effect of material properties

The yield stress $\tau_y$ describes the stress above which the liquid begins to flow. The flow consistency index $K$ sets the viscosity scale of the liquid, while the flow-behavior index $n$ describes its shear-rate dependence. Values $0 < n < 1$ correspond to shear-thinning liquids, for which the apparent viscosity decreases with increasing shear rate; $n = 1$ corresponds to Newtonian behavior; and $n > 1$ corresponds to shear-thickening behavior.

Extensional rheometer tests of the termite secretion yielded an apparent extensional viscosity of approximately $\eta_E \sim 100$ Pa $\cdot$ s, with pronounced hardening at larger strains. Direct steady-shear measurements were not available because of the small secretion volumes and the rapid evolution of the material after ejection. As an order-of-magnitude estimate of the low-strain viscosity scale, we assume that the initial response can be approximated as Newtonian and use the Trouton relation

$$T_r = \frac{\eta_E}{\eta}.$$

For a Newtonian fluid, $\eta_E = 3\eta$, such that $T_r = 3$. Applying this relation to the initial extensional viscosity gives a characteristic shear-viscosity scale of $\eta \sim 33$ Pa $\cdot$ s, which we use as a starting value for the flow model.

At strains $\varepsilon > 2$, the apparent Trouton ratio increases and reaches a plateau of approximately $T_r = 10.5$, indicating an increasingly non-Newtonian extensional response. For non-Newtonian fluids, however, extensional and shear viscosities cannot be related directly without additional constitutive information because the response depends on deformation type, deformation rate, and deformation history. The extensional measurements therefore do not by themselves determine whether the secretion is shear-thinning or shear-thickening.

We consequently explored shear-thinning, Newtonian, and shear-thickening behavior parametrically. The initial flow-behavior indices were $n = 1$ for the Newtonian case, $n = 0.2$ for a strongly shear-thinning liquid, and $n = 1.8$ for a shear-thickening liquid. These cases were used to evaluate how the assumed shear-rate dependence affects the pressure required for extrusion.

Fig. S7b shows the pressure drop for each shape function. The ordering is the same for all three constitutive cases: the linear nozzle requires the lowest pressure and the constant-diameter nozzle the highest. The required pressures span several orders of magnitude, and only the shear-thinning case yields values consistent with the pressure range estimated for ejection (main text). This result supports the use of a shear-thinning constitutive model for the subsequent nozzle comparisons, although it does not constitute a direct measurement of the natural secretion's steady-shear response. Table S6 lists the values and Fig. S7c compares the shear-thinning case alone. The termite nozzle shows a higher pressure drop than the linear nozzle but a lower one than all other nozzles.

A sweep of the flow-behavior index for shear-thinning liquids confirms the exponential growth of the required

TABLE S3. **Coefficients defining the comparison nozzle shape functions.** All comparison geometries were constrained by the nozzle length $L$ and inlet and outlet diameters $D_{\text{inlet}}$ and $D_{\text{outlet}}$, respectively.

| **Shape** | $\boldsymbol{a}$ | $\boldsymbol{b}$ | $\boldsymbol{c}$ | $\boldsymbol{d}$ |
|---|---|---|---|---|
| Termite nasus | $-20.727$ | 30.255 | $-0.311$ | — |
| Constant diameter | $D_{\text{outlet}}$ | — | — | — |
| Linear | $D_{\text{inlet}}$ | $\dfrac{D_{\text{outlet}} - D_{\text{inlet}}}{L}$ | — | — |
| Quadratic | $D_{\text{inlet}}$ | $\dfrac{2(D_{\text{outlet}} - D_{\text{inlet}})}{L}$ | $\dfrac{D_{\text{inlet}} - D_{\text{outlet}}}{L^2}$ | — |
| Cubic | $D_{\text{inlet}}$ | $\dfrac{3(D_{\text{outlet}} - D_{\text{inlet}})}{L}$ | $\dfrac{3(D_{\text{inlet}} - D_{\text{outlet}})}{L^2}$ | $\dfrac{D_{\text{outlet}} - D_{\text{inlet}}}{L^3}$ |
| Exponential | $D_{\text{inlet}}$ | $\dfrac{\ln(D_{\text{outlet}}/D_{\text{inlet}})}{L}$ | — | — |

TABLE S4. **Numerical coefficients used for the nozzle shape functions.**

| **Shape** | $\boldsymbol{a}$ | $\boldsymbol{b}$ | $\boldsymbol{c}$ | $\boldsymbol{d}$ |
|---|---|---|---|---|
| Termite nasus | $-20.727$ | 30.255 | $-0.311$ | — |
| Constant diameter | 0.032 | — | — | — |
| Linear | 0.173 | $-0.104$ | — | — |
| Quadratic | 0.173 | $-0.208$ | 0.077 | — |
| Cubic | 0.173 | $-0.312$ | 0.231 | $-0.057$ |
| Exponential | 0.173 | $-1.247$ | — | — |

TABLE S5. **Rheological parameters used in the Herschel–Bulkley parametric study.**

| **Flow behavior** | $\boldsymbol{\tau_y}$ (Pa) | $\boldsymbol{k}$ (Pa s$^n$) | $\boldsymbol{n}$ |
|---|---|---|---|
| Newtonian | 0 | 33 | 1.0 |
| Shear-thinning | 0 | 33 | 0.2 |
| Shear-thickening | 0 | 33 | 1.8 |

TABLE S6. **Pressure drop predicted for each nozzle geometry as a function of flow-behavior index** $n$**.**

| **Nozzle shape** | $\boldsymbol{n = 0.2}$ | $\boldsymbol{n = 1.0}$ | $\boldsymbol{n = 1.8}$ |
|---|---|---|---|
| Termite | $1.04 \times 10^4$ | $7.61 \times 10^6$ | $9.38 \times 10^9$ |
| Constant | $3.84 \times 10^4$ | $8.55 \times 10^7$ | $1.90 \times 10^{11}$ |
| Linear | $9.24 \times 10^3$ | $6.41 \times 10^6$ | $7.98 \times 10^9$ |
| Exponential | $1.33 \times 10^4$ | $1.26 \times 10^7$ | $1.76 \times 10^{10}$ |
| Quadratic | $1.59 \times 10^4$ | $2.00 \times 10^7$ | $3.37 \times 10^{10}$ |
| Cubic | $2.04 \times 10^4$ | $3.11 \times 10^7$ | $5.78 \times 10^{10}$ |

Pressure drop is reported in Pa.

pressure with $n$ (Fig. S7d); the curves remain parallel, showing that the ranking of the shapes is preserved across a wide range of $n$.

The pressure drop response is visually apparent across all of the nozzle shape functions, with the linear and termite nozzles each experiencing max pressures of ∼2E5 Pa, while the cubic shape function experiences a max pressure of about ∼5E5, dramatically affecting the corresponding shear rates at the walls and dynamic viscosity of the shear-thinning fluid (Fig. S8).

#### F. Flow regime in the natural and scaled nozzles

For the non-Newtonian secretion considered here, the Reynolds number is not uniquely defined because the apparent viscosity depends on shear rate. We therefore use an outlet-based effective-viscosity estimate to characterize the approximate flow regime. For a power-law fluid,

$$\tau = k\dot{\gamma}^{\,n},$$

the apparent viscosity is

$$\mu_{\text{eff}} = k\dot{\gamma}^{\,n-1}.$$

For fully developed flow in a circular tube, the wall shear rate of a power-law fluid can be estimated using the Rabinowitsch correction,

$$\dot{\gamma}_w \approx \frac{8u}{D_{\text{outlet}}}\left(\frac{3n+1}{4n}\right),$$

where $u$ is the mean outlet velocity and $D_{\text{outlet}}$ is the outlet diameter. Using the experimentally estimated outlet velocity $u \sim 0.4$ m/s, $D_{\text{outlet}} \sim 32$ $\mu$m, $k = 33$ Pa s$^n$, $n = 0.46$, and $\rho \sim 900$ kg/m$^3$, this gives an effective-viscosity Reynolds number

$$Re_{\text{eff}} = \frac{\rho u D_{\text{outlet}}}{\mu_{\text{eff}}} \sim 0.2.$$

This estimate is intended only as an order-of-magnitude characterization because the nasus is tapered and the secretion is more accurately described as a yield-stress, shear-thinning material. Nevertheless, $Re_{\text{eff}} < 1$ indicates that viscous stresses dominate inertia near the nozzle outlet.

To enable fabrication and controlled testing, the termite-inspired nozzle was enlarged by a factor of 20, yielding an outlet diameter of $D_{\text{outlet}} = 0.64$ mm. Exact dynamic similarity with the natural system was not imposed because changing the geometric scale alters the relationships among velocity, flow rate, shear rate, and the apparent viscosity of the non-Newtonian fluid. Instead, the scaled experiments preserved the principal nozzle geometry while allowing pressure and nozzle shape to be varied independently. For the synthetic ejection tests, with $u \sim 0.64$ m/s, $D_{\mathrm{outlet}} = 0.64$ mm, $k \approx 29.5$ Pa s$^{\mathrm{n}}$, $n \approx 0.44$, and $\rho \sim 1050$ kg/m$^3$, the same estimate gives $Re_{\mathrm{eff}} \approx 3$, so inertial and viscous stresses are comparable at the outlet and the flow remains laminar.

Flow rates through the fabricated nozzle profiles were measured under controlled applied pressures and compared with the trends predicted by the numerical model. These experiments therefore provide an experimental comparison of hydraulic performance among nozzle geometries rather than a dynamically scaled replica of natural termite ejection. The larger oscillatory-ejection experiments described separately likewise constitute a synthetic design-space study inspired by termite behavior rather than a quantitative model of the natural system.

#### G. Rheology of the synthetic secretion analog

The shear rheology of the hydrogel:water mixture (30:70 by weight) was measured in a shear-rate sweep on a rheometer. A Herschel–Bulkley fit gave $\tau_y \approx 609$ Pa, $k \approx 29.5$ Pa s$^{\mathrm{n}}$, and $n \approx 0.44$, close to the parameters inferred for the termite secretion ($\tau_y = 298$ Pa, $k = 33$ Pa s$^{\mathrm{n}}$, $n = 0.46$). The viscosity versus shear rate and the storage and loss moduli show the shear-thinning character of the soil:hydrogel inks relative to the hydrogel:water mixture (Fig. S9).

##### 1. *Ejection of the synthetic analog through the scaled nozzle*

Ejection velocity was measured from the high-speed video by identifying the frame in which the filament tip first left the nozzle, selecting a frame $n$ frames later, and measuring the filament length $L$ from nozzle tip to filament tip in that frame in ImageJ. With camera frame rate $f_{\mathrm{cam}}$, the elapsed time is $\Delta t = n/f_{\mathrm{cam}}$ and the mean tip velocity is

$$v = \frac{L}{\Delta t} = \frac{L\, f_{\mathrm{cam}}}{n}.$$

Measured near the nozzle exit at the onset of ejection, $v = 640$ and 650 mm/s at 45 and 50 psi (310 and 345 kPa), respectively. At 40 psi (276 kPa), the velocity measured later in the pulse, once flow was fully developed, reached 2000 mm/s (Fig. S10).

##### 2. *Extrusion printing with the termite-shaped nozzle*

The termite-shaped nozzle is well suited to 3D printing of non-Newtonian materials, since the wall shear stress near the outlet lowers the apparent viscosity and hence the pressure required for a given flow rate. Direct ink writing (DIW) nozzles often taper in the same way to reduce the pressure drop [14]. Soil-loaded synthetic-analog mixtures can be used as a printable ink, as demonstrated by printed termite mound structures (Fig. S14).

Samples printed with varying soil:hydrogel compositions and dried at varying temperatures were subjected to compression tests (Fig. S14a–c); the strongest combination was the 1:5 mix dried at 70 °C (Fig. S14c). Fig. S14d shows a termite-inspired mound printed with this material, during printing and after completion.

Nozzles were modeled in SolidWorks, sliced with Lychee Slicer, and printed on a digital light processing printer (Sonic Mini 8K, Phrozen) with Aqua-Gray 8K resin (Phrozen) at a 50 $\mu$m layer height and 1.8 s exposure time. Termite mounds were printed with the 50× termite-shaped nozzle (1.6 mm outlet diameter) from a 30 mL syringe at 11 psi (76 kPa) and a 30 mm/s feed rate, using the 1:5 soil:hydrogel mixture.

### V. Supplementary videos

**Supplementary video S1. Defensive ejection and head scanning in *Nasutitermes corniger*.** High-speed footage (5000 fps) from Barro Colorado Island and Gamboa, Panama, of soldiers ejecting secretion toward a trap-jaw ant, the lateral head scan during ejection, and a trap-jaw ant attempting to groom the secretion from its body. Scale bars, 1 mm. Playback speed as indicated.

**Supplementary video** S2. Liquid rope coiling, lasso deposition, and extensional rheology. Rope coiling of the ejected filament on a glass slide (*Nasutitermes* sp., 10 000 fps); deposition of a liquid lasso during head scanning (5000 and 2500 fps); and capillary thinning of fresh secretion in the field extensional rheometer for *Nasutitermes* sp. (Peru) and *N. corniger* (Panama), including buckling of the arrested *N. corniger* filament when the upper plate is lowered. Scale bars, 1 mm and 500 $\mu$m as indicated.

**Supplementary video** S3. Nasus morphology and scaled synthetic ejection. Micro-CT reconstruction of the nasus (*Nasutitermes* sp.); ejection of the synthetic secretion analog through a twenty-fold enlarged termite-shaped nozzle at 640 and 650 mm/s (18 600 fps); oscillatory ejection producing a synthetic liquid lasso; and deposition of the synthetic lasso onto a model trap-jaw ant. Scale bars, 1 and 2 mm as indicated.

### VI. Supplementary figures

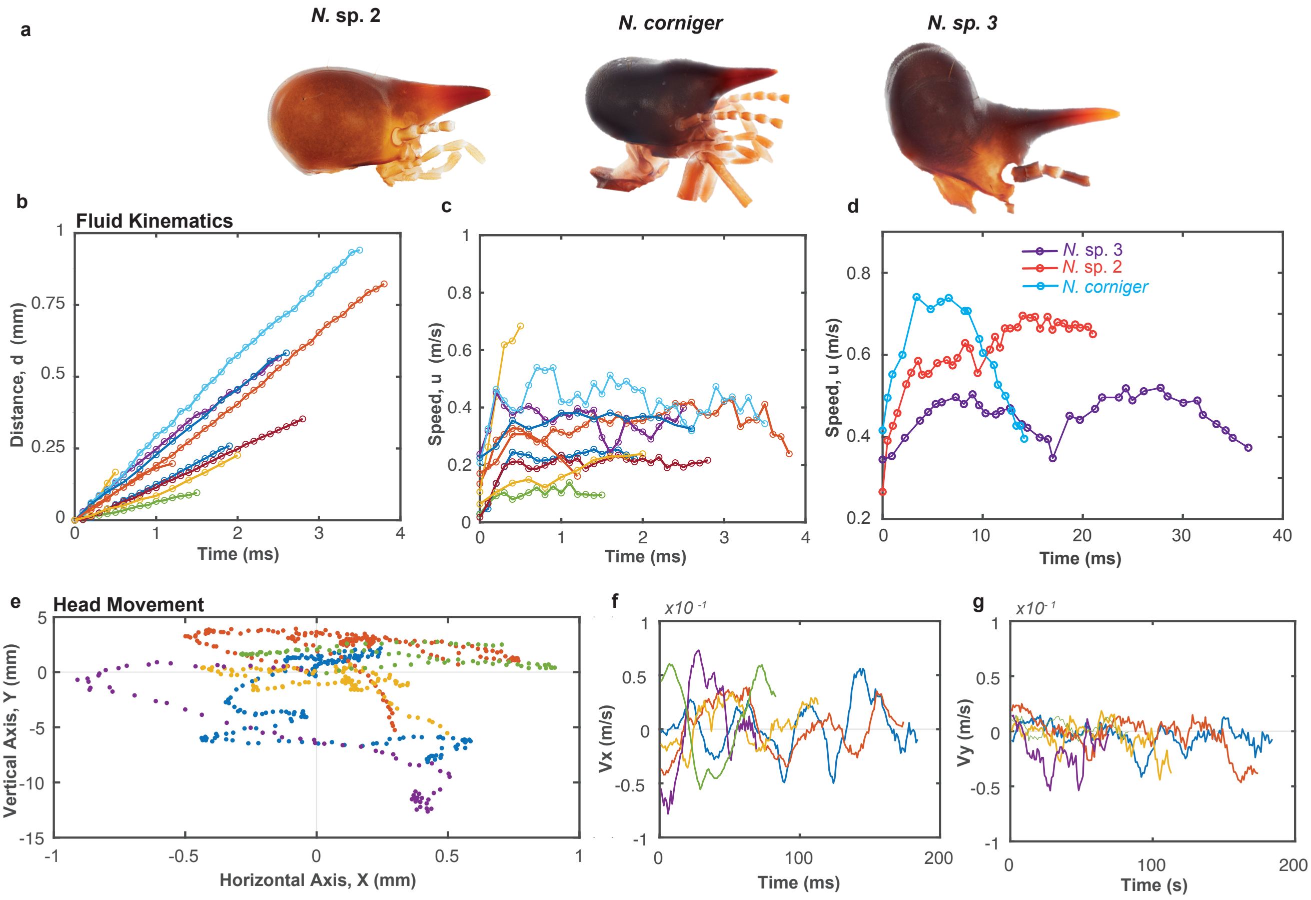


Fig. S1. **Ejection kinematics and head scanning. a.** Nasus morphologies of cone-headed Nasutitermitinae soldiers. **b.** Tracking of jet displacement as the secretion exits the nasus prior to substrate impact (Nasutitermitinae). **c.** Jet speed versus time for Nasutitermitinae soldiers. **d.** Comparison of jet speeds across three Nasutitermitinae species. **e.** Head scanning during ejection, showing an initial horizontal sweep followed by retreat. **f–g. f,g.** Head velocity during ejection in the horizontal direction (perpendicular to the initial stance) and the vertical direction (parallel to the initial stance), respectively. Frame rates: 10,000 fps (b–d) and 5,000 fps (e–g)

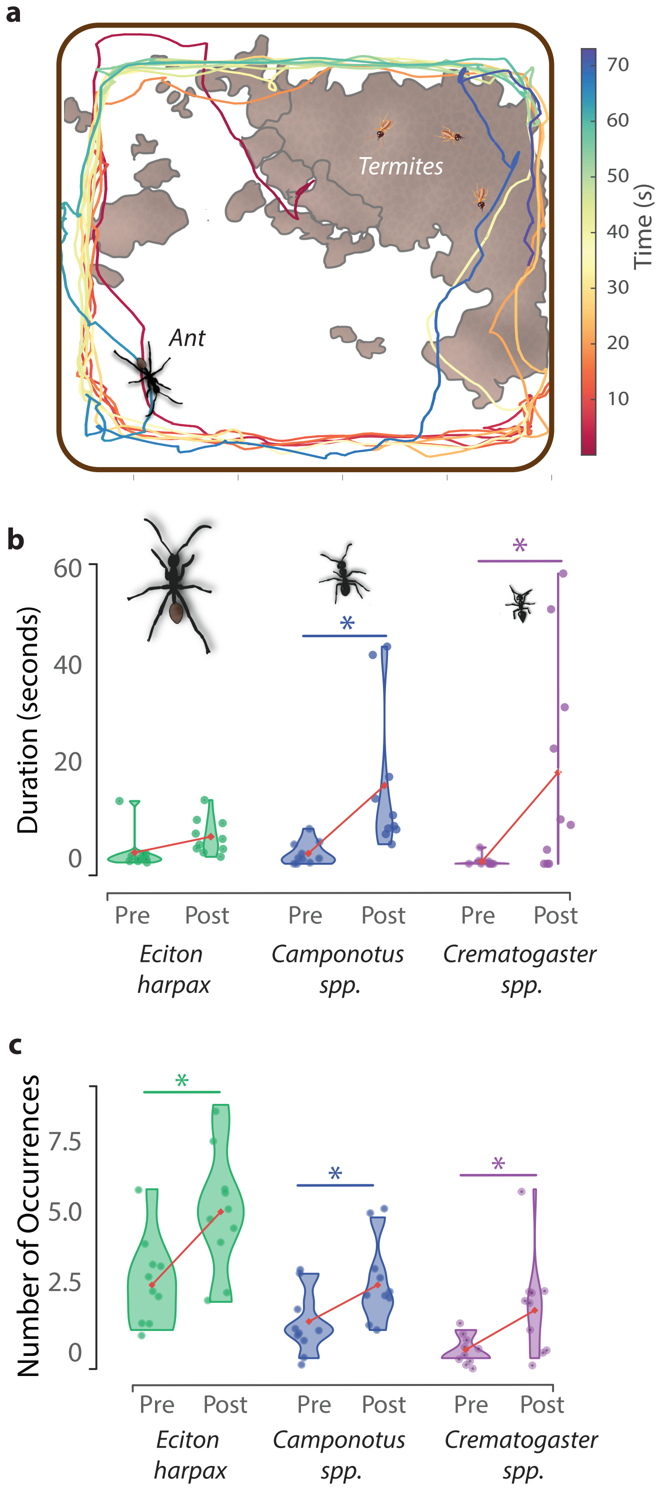


Fig. S2. **Termite exposure increases grooming behavior in ant predators.** **a.** Representative ant trajectories during encounters with soldier termites. Colored tracks indicate time progression; ants move primarily along the arena perimeter, whereas termites remain clustered near the nest material. **b.** Fraction of time spent grooming before (Pre) and after (Post) the encounter for *Eciton harpax*, *Camponotus* sp., and *Crematogaster* sp. **c.** Number of grooming bouts before and after the encounter for the same taxa. Violin plots show the data distributions, points represent individual ants ($n = 10$ exposed and 3 control per taxon), lines connect group means, and asterisks indicate significant post-hoc contrasts.

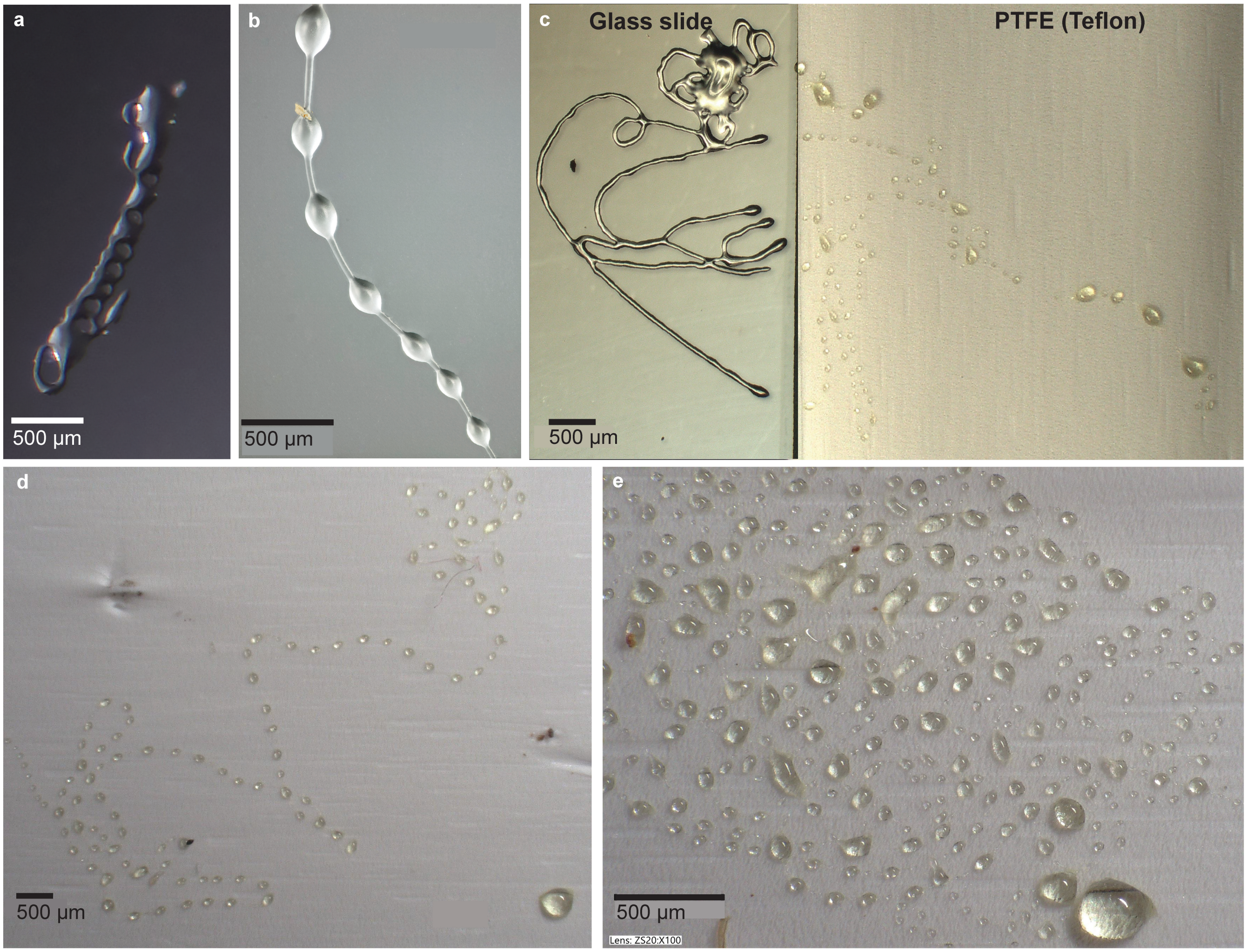


Fig. S3. **Deposited secretion on glass and PTFE. a.** Remnants of termite secretion on a glass slide showing overlapping coils. **b.** Bead-on-a-string (BOAS) morphology emerging from termite secretion on glass, indicating the viscoelastic properties of the fluid (*Nasutitermes* sp.). **c.** Termite secretion collected on glass and PTFE surfaces from the same secretion event, illustrating the effect of surface wettability on fluid spreading and BOAS formation (*Nasutitermes corniger*). **d.** Zoomed view of BOAS formation on glass. **e.** Zoomed view of termite secretion on PTFE showing pronounced beading (*Nasutitermes corniger*).

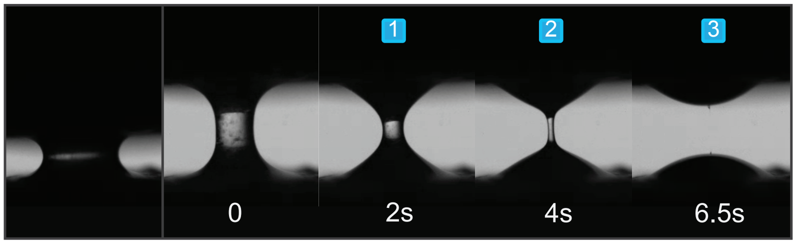


Fig. S4. **Complete filament breakup of *Nasutitermes* sp. secretion.** Capillary-thinning sequence from the field extensional rheometer (500 fps) showing complete breakup with no persistent filament, in contrast to *N. corniger* (main text Fig. 4).

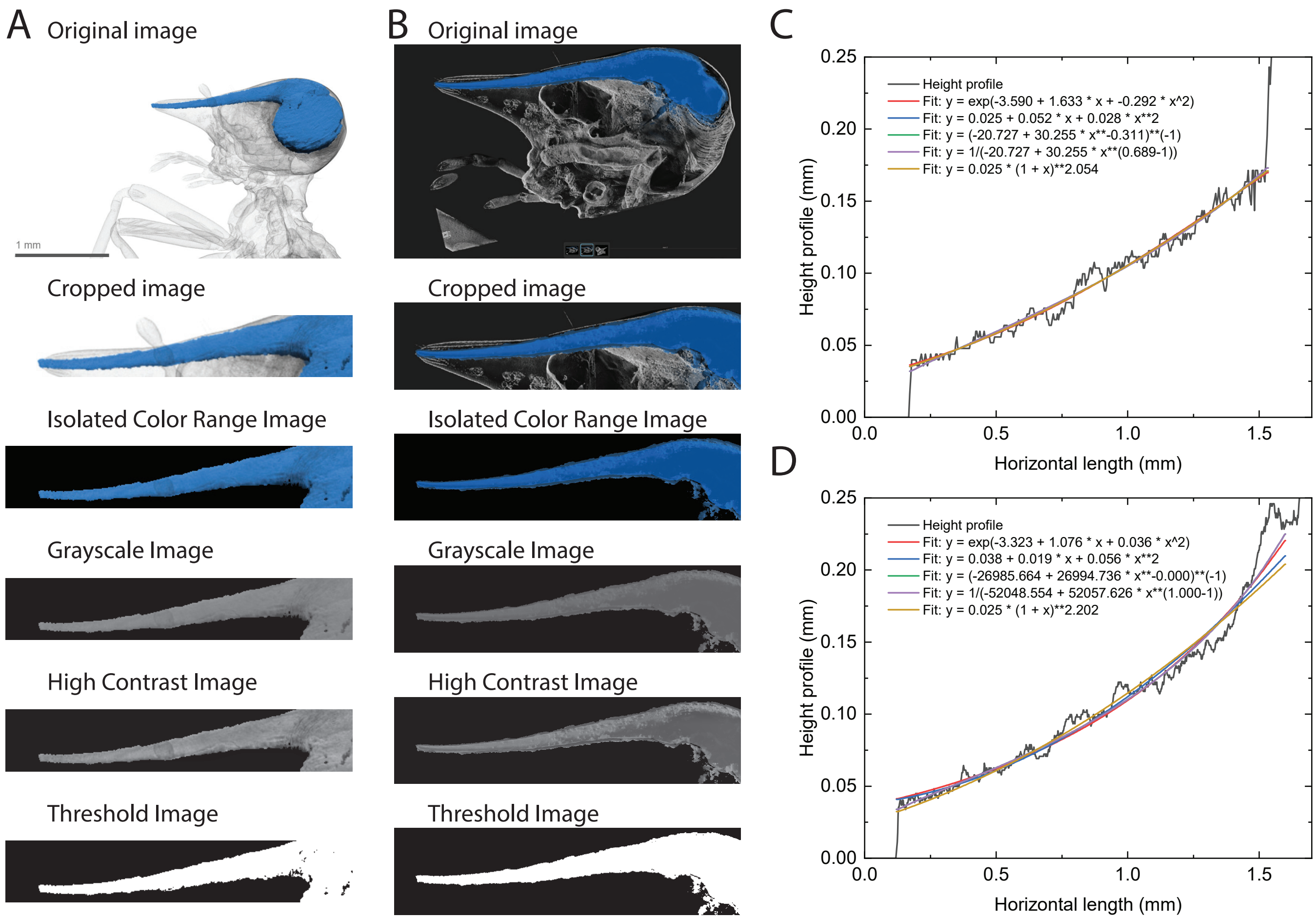


Fig. S5. **Termite nozzle profile extraction from micro-CT. a.** Termite head 1 with the extraction steps 1–7 described in the text. **b.** Termite head 2 with the same steps. **c.** Height profile and fitted functions for head 1. **d.** Height profile and fitted functions for head 2.

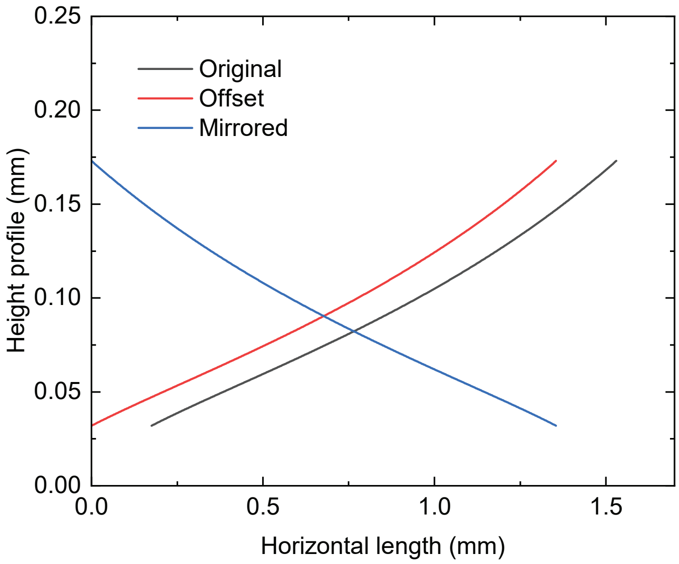


Fig. S6. **Termite nozzle shape transformations.** Original, offset, and mirrored forms of the fitted nasus profile used to place the inlet at $x = 0$ in the analytical and numerical models (Section IV).

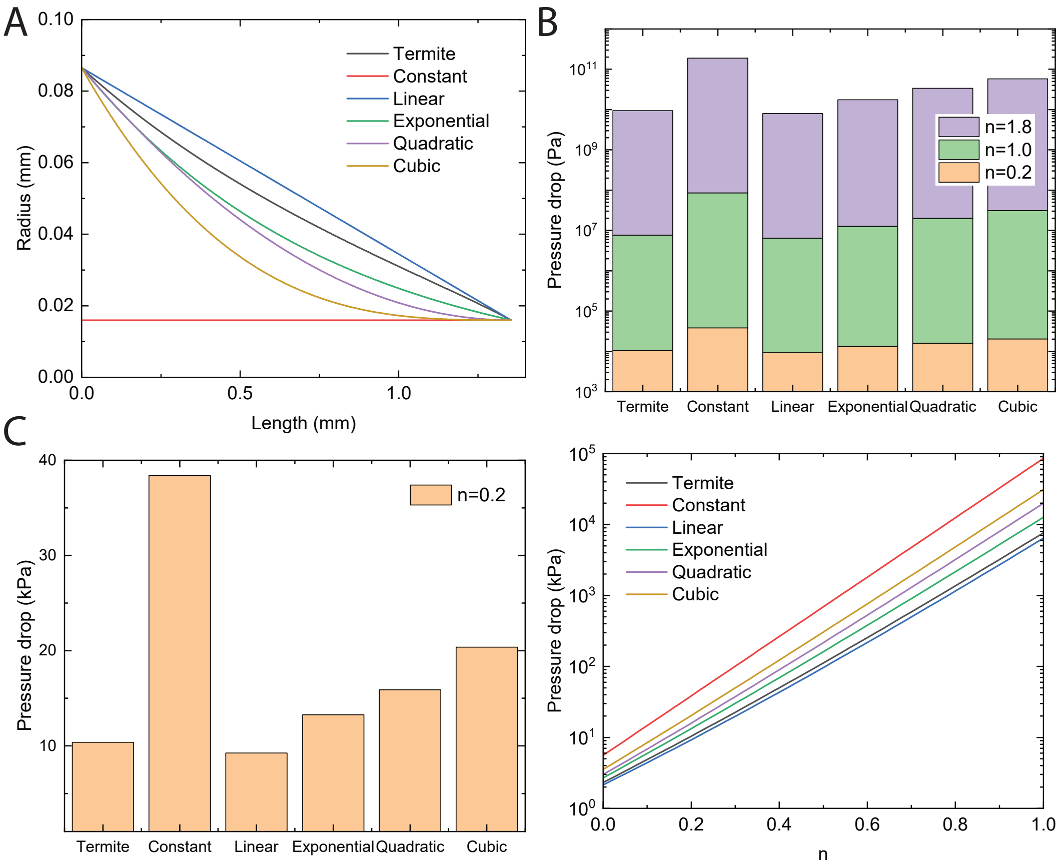


Fig. S7. **Pressure drop simulation for each nozzle profile.** **a.** Shape function of each studied profile. **b.** Pressure drop for each shape function for the three constitutive cases of Table S5. **c.** Pressure drop for each shape function for the shear-thinning case ($n = 0.2$). **d.** Pressure drop for each shape function over varying values of $n$.

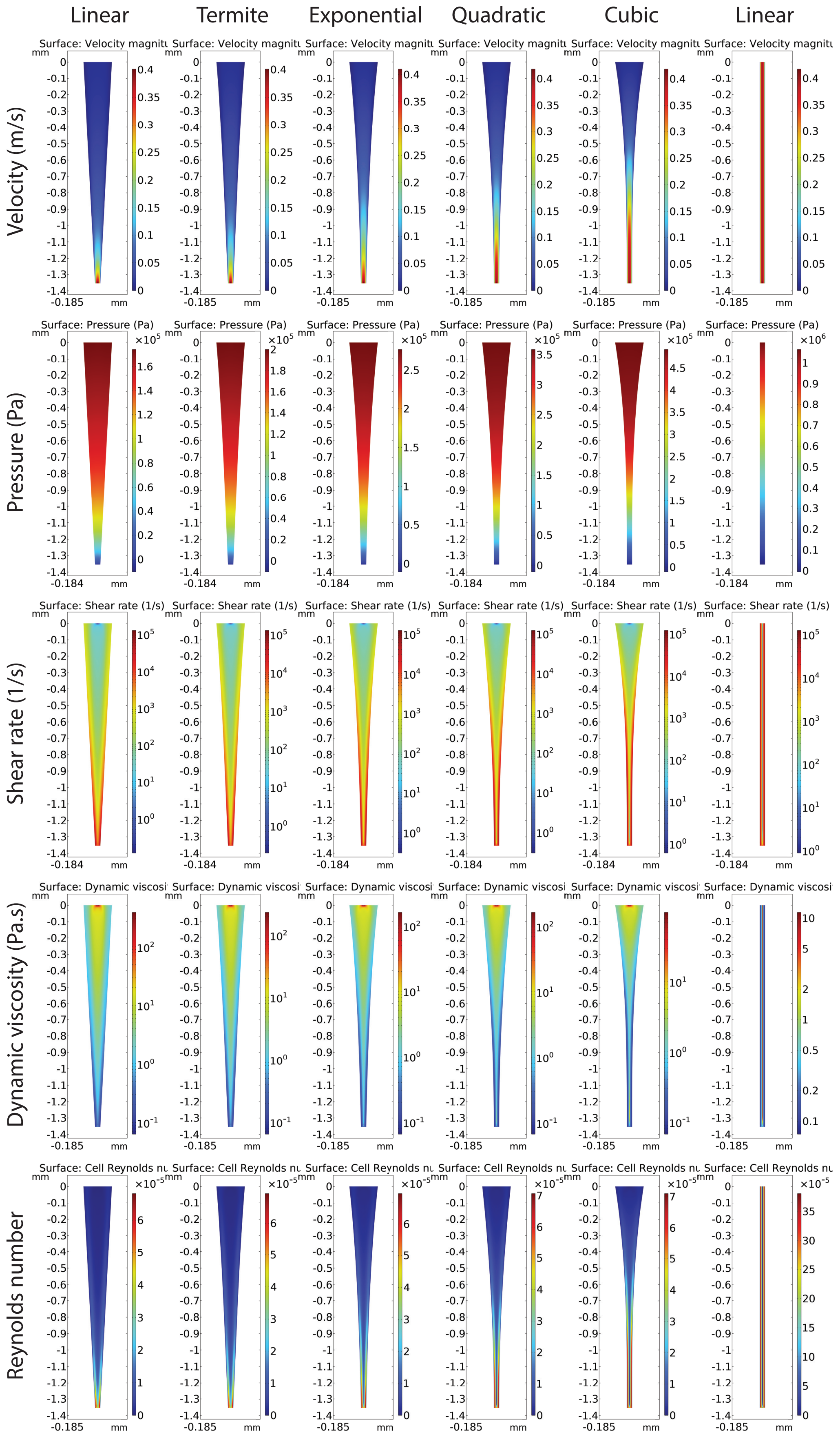


Fig. S8. **Simulated flow through each nozzle type.** Under an applied pressure, the velocity, pressure, shear rate, dynamic viscosity, and Reynolds number response for the linear, termite, exponential, quadratic, cubic, and linear shape functions.

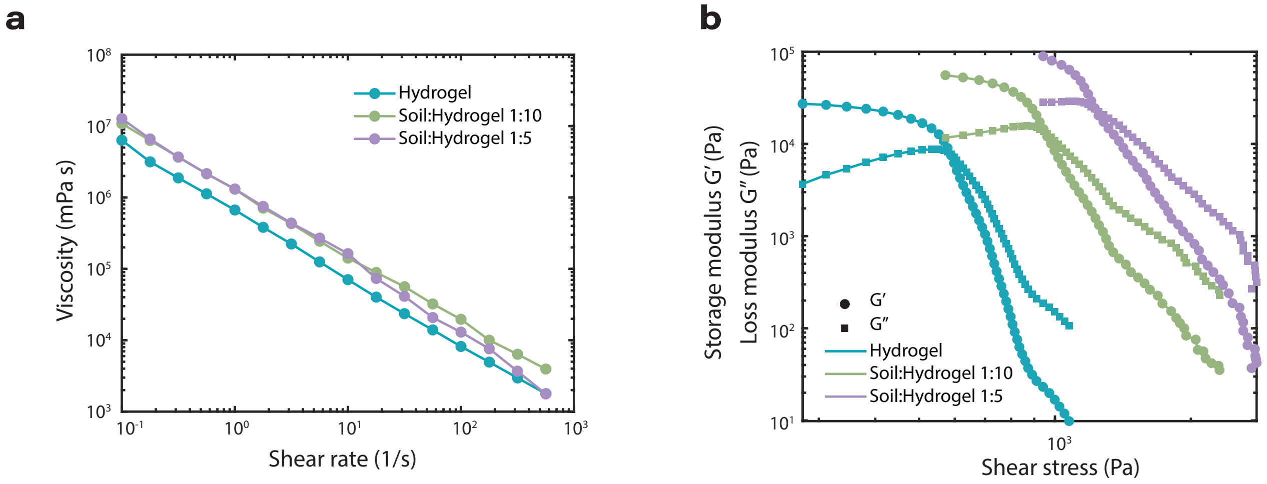


Fig. S9. **Rheology of inks. a.** Viscosity versus shear rate for the hydrogel mixtures. **b.** Storage ($G'$) and loss ($G''$) moduli for the hydrogel mixtures.

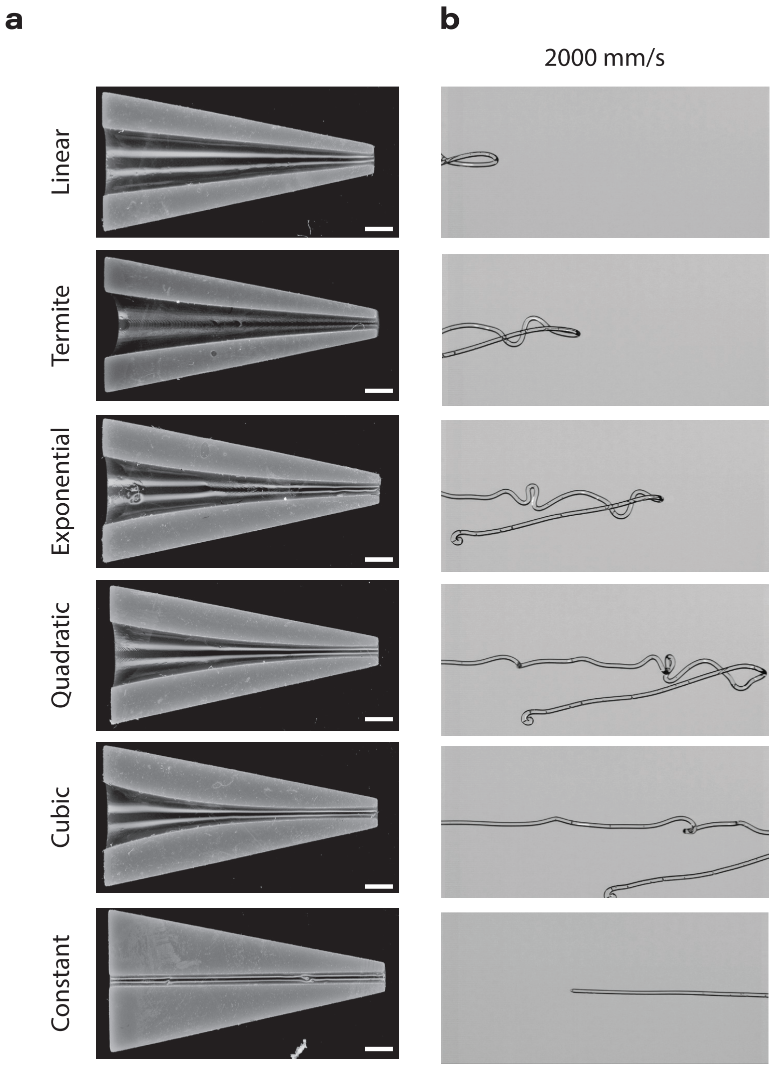


Fig. S10. **Fabricated nozzles and synthetic ejection. a.** 3D-printed nozzle cross-sections for each shape function. **b**, High-speed images (18,600 fps) of fully developed hydrogel flow ejected from 20× termite nozzle under an applied pressure of 40 psi (276 kPa). Scale bars are 1 mm.

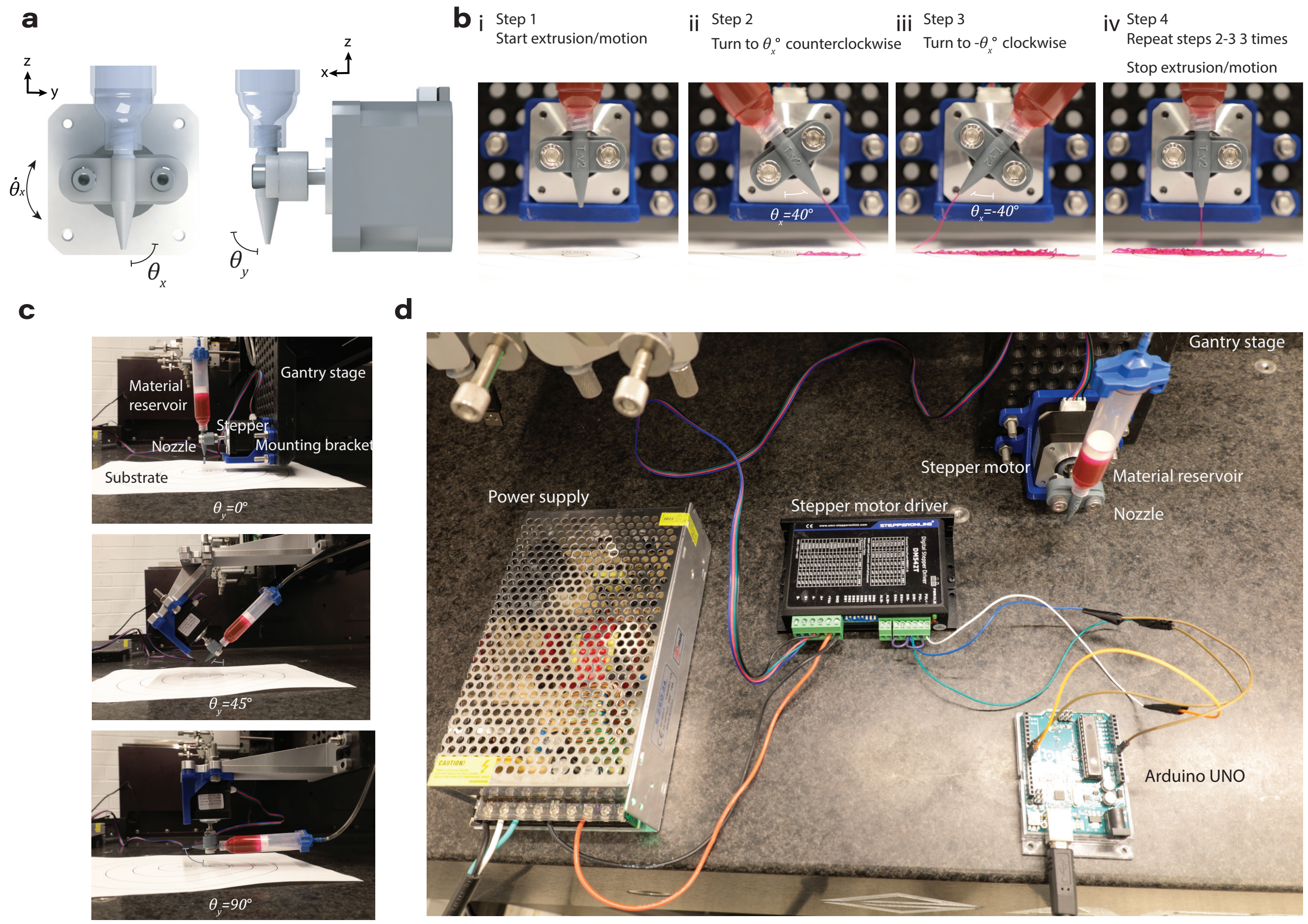


Fig. S11. **Mechanical liquid lasso test setup.** **a.** Definitions of the angular velocity ($\dot{\theta}_x$), pitch angle ($\theta_y$), and yaw angle ($\theta_x$). **b.** Motion of nozzle during oscillation. **c.** Mounting setups for each level of pitch angle $\theta_y$. **d.** Test setup components for stepper motor control.

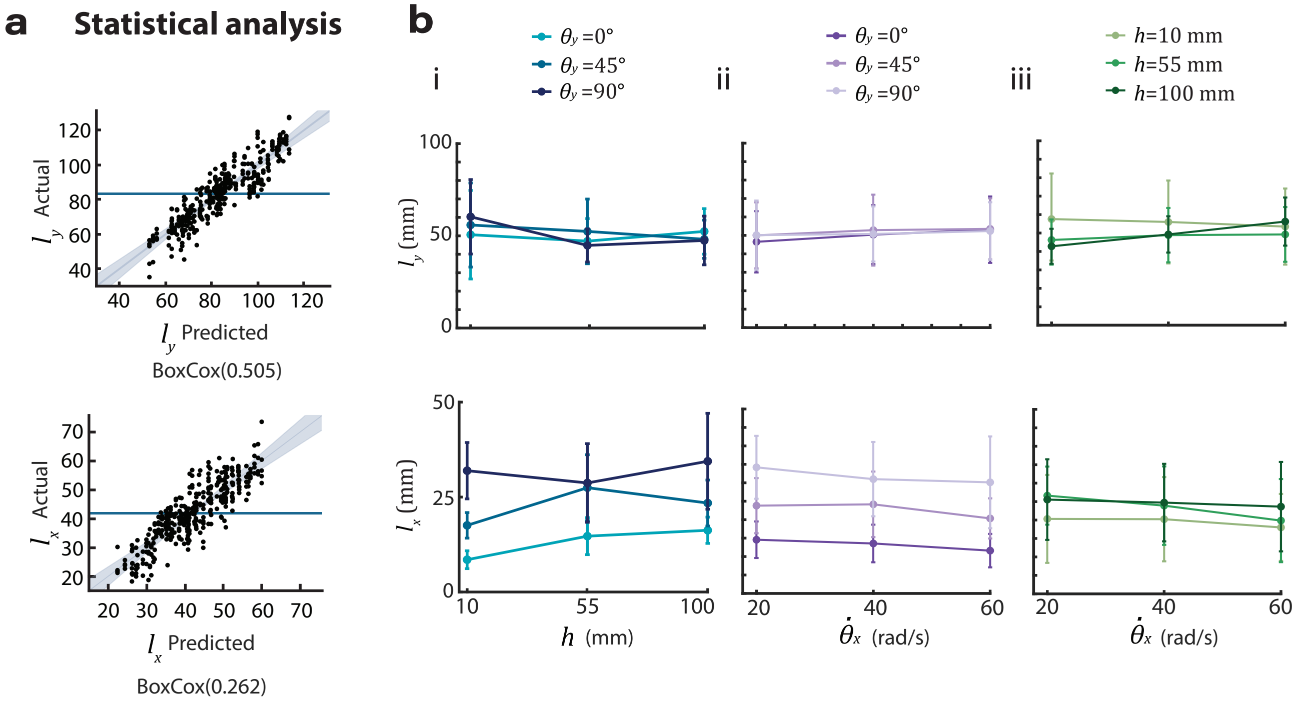


Fig. S12. **Extended statistical analysis results. a.** Model fits for width ($l_y$) and depth ($l_x$) after Box-Cox transformation. **b.** Interaction effects for i) yaw angle ($\theta_x$) and nozzle height ($h$), ii) pitch angle ($\theta_y$) and angular velocity ($\dot{\theta}_x$), and iii) nozzle height ($h$) and angular velocity ($\dot{\theta}_x$).

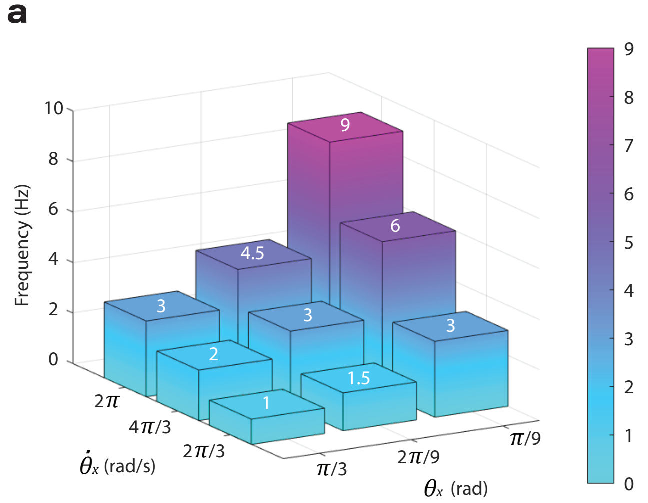


Fig. S13. **Oscillation frequency of the mechanical nozzle.** Nozzle oscillation frequency as a function of $\theta_x$ and $\dot{\theta}_x$.

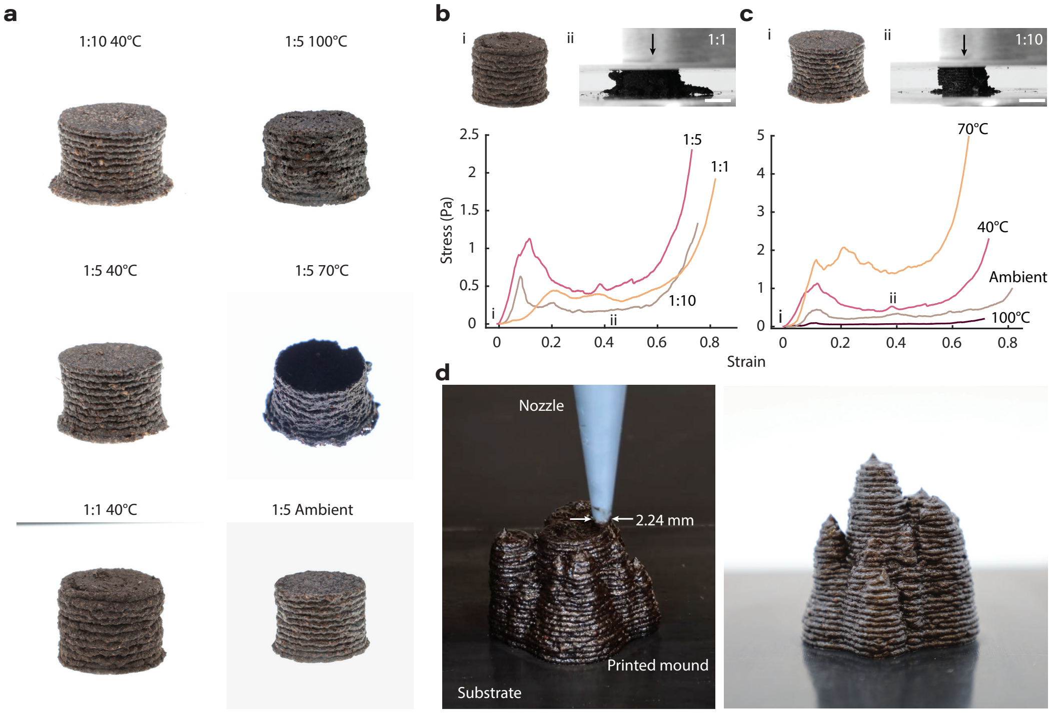


Fig. S14. **3D printing with the synthetic secretion analog. a.**, Compression test samples for each soil:hydrogel composition at varying drying temperatures (shrinkage occurs during drying as water evaporates). **b.** Compression test results of printed soil:hydrogel samples, varying the composition ratio at 40°C drying temperature. i) 1:1 ratio printed sample. ii) 1:1 ratio sample crumbling under compression. **c.** Compression test results of printed 1:5 soil-hydrogel samples, varying the drying temperature from ambient to 100°C. i) 1:10 ratio printed sample. ii) 1:10 ratio sample squishing under compression. **d.** 3D-printed bio-inspired termite mounds with the soil-hydrogel material. Scale bars are 7 mm.

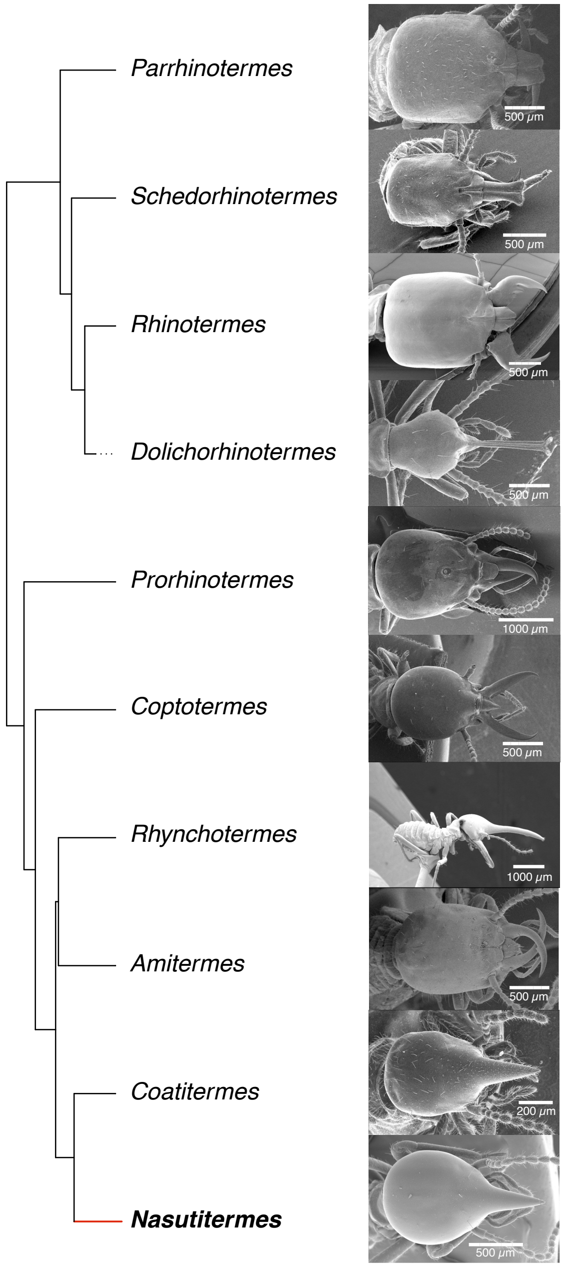


Fig. S15. **Comparative head shape across Isoptera.** Pruned phylogeny of termite genera highlighting groups with pores or nozzles, presented alongside a scanning electron microscope image of the dorsal view of a termite head. The focal genus of this study, *Nasutitermes*, is shown in bold to indicate its placement within the clade.

[6] P. A. Latreille, *Histoire naturelle, générale et particulière, des crustacés et des insectes*, Vol. 3 (F. Dufart, Paris, 1802) original description of Formica apicalis on p. 204.

[7] C. Emery, Voyage de m. ch. alluaud dans le territoire d'assinie. formicides, Annales de la Société entomologique de France **60**, 553 (1892), journal volume may carry the year 1891; nomenclatural publication year is 1892.

[8] T. Eisner, I. Kriston, and D. J. Aneshansley, Defensive behavior of a termite (nasutitermes exitiosus), Behavioral Ecology and Sociobiology **1**, 83 (1976).

[9] C. L. Yaws, Chemical properties handbook: Physical, thermodynamic, environmental, transport, safety and health related (1999).

[10] U. C. Guard, *Chemical Hazards Response Information System (CHRIS) Hazardous Chemical Data*, Tech. Rep. (Washington, D.C., 1984) 1984-5.

[11] G. H. McKinley, Visco-elasto-capillary thinning and break-up of complex fluids, Rheology Reviews **2005**, 1 (2005).

[12] Originlab, corporation - data analysis and graphing software - 2d graphs, 3d graphs, contour plots, statistical charts, data exploration, statistics, curve fitting, signal processing, and peak analysis.

[13] I. Langmuir, The adsorption of gases on plane surfaces of glass, mica and platinum., Journal of the American Chemical Society **40**, 1361 (1918).

[14] T. Schuller, M. Jalaal, P. Fanzio, and F. J. Galindo-Rosales, Optimal shape design of printing nozzles for extrusion-based additive manufacturing, Additive Manufacturing **84**, 104130 (2024).
